# DeepSP: Deep Learning-Based Spatial Properties to Predict Monoclonal Antibody Stability

**DOI:** 10.1101/2024.02.28.582582

**Authors:** Lateefat Kalejaye, I-En Wu, Taylor Terry, Pin-Kuang Lai

## Abstract

Therapeutic antibody development, manufacturing, and administration face challenges due to high viscosities and aggregation tendencies often observed in highly concentrated antibody solutions. This poses a particular problem for subcutaneous administration, which requires low-volume and high-concentration formulations. The spatial charge map (SCM (mAbs, 8 (1) (2015), pp. 43-48)) and spatial aggregation propensity (SAP (PNAS. 2009; 106:11937–42) are two computational techniques proposed from previous studies to aid in predicting viscosity and aggregation, respectively. These methods rely on structural data derived from molecular dynamics (MD) simulations, which are known to be time-consuming and computationally demanding. DeepSCM (CSBJ. 2022, 20:2143-2152), a deep learning surrogate model to predict SCM scores in the entire variable region, was used to screen high-concentration antibody viscosity. DeepSCM is solely based on sequence information, which facilitates high throughput screening. This study further utilized a dataset of 20,530 antibody sequences to train a convolutional neural network deep learning surrogate model called Deep Spatial Properties (DeepSP). DeepSP directly predicts SAP and SCM scores in different domains of antibody variable regions based solely on their sequences without performing MD simulations. The linear correlation coefficient (R) between DeepSP scores and MD-derived scores for 30 properties achieved values between 0.76 and 0.96 with an average of 0.87 on the test set (N=2053). DeepSP was employed as features to build machine learning models to predict the aggregation rate of 21 antibodies. We observed remarkable results with R = 0.97 and a mean squared error (MSE) of 0.03 between the experimental and predicted aggregation rates, leave-one-out cross-validation (LOOCV) yielded R = 0.75 and MSE = 0.18, which is similar to the results obtained from the previous study using MD simulations. This result demonstrates that the DeepSP approach significantly reduces the computational time required compared to MD simulations. The DeepSP model enables the rapid generation of 30 structural properties that can also be used as features in other research to train machine learning models for predicting various antibody properties, such as viscosity, aggregation, or other properties that can influence their stability, using sequences only. The code and parameters are freely available at https://github.com/Lailabcode/DeepSP

**Highlights:**

- Deep learning applied to develop a surrogate model (DeepSP) to rapidly predict 30 spatial properties of monoclonal antibodies that are usually calculated from MD simulations, using only sequences.
- The DeepSP models achieved a linear correlation ranging between 0.76 and 0.96 with an average of 0.87, between the actual (MD simulation) and predicted score for all properties.
- DeepSP features were employed to build a model to predict aggregation rates of antibodies obtained from a previous study. A strong correlation of 0.97, and LOOCV correlation of 0.75 were achieved between the actual and predicted aggregation rates.
- DeepSP can be employed to generate antibody-specific features that can be used to train different machine learning models to predict antibody stability.

## 1. Introduction

Highly concentrated antibody solutions often exhibit high viscosities^1^, aggregation tendencies^2,3^, and various forms of instability, posing significant challenges in antibody-drug development, manufacturing, and administration. Subcutaneous administration requires low-volume and high-concentration formulations^4–7^. With the increasing desire to improve patient convenience and compliance with monoclonal antibodies (mAbs) by moving away from intravenous and towards subcutaneous mode of administration^8,9^, solutions must be developed to overcome the challenges faced when formulating highly concentrated antibody drugs. The antibody sequence is critical for antibody engineering^10^ and acts as a key determinant for high viscosity^1^, and other instability issues of highly concentrated solutions. Therefore, developing a sequence-based model that can be used to identify problematic antibodies is desired.

Agrawal et al.^1^ developed the spatial charge map (SCM) as a computational tool via molecular dynamics (MD) simulation that can be used for antibody screening to effectively differentiate low or high viscosity antibodies. Naresh et al.^2^ developed the spatial aggregation propensity (SAP) as a computational tool via MD simulation that can be used to identify the location and size of aggregation-prone regions and allows target mutations of those regions to engineer antibodies for improving stability. In addition, coarse-grained (CG) models have been implemented in different studies^11–15^ to help screen antibody viscosity and other developability issues. However, these methods are computationally costly and require structural information, which is a significant application bottleneck.

In recent years, machine learning techniques have been adopted in predicting high concentration antibody stability. Lai et al.^16^ used machine learning to determine the molecular descriptors responsible for the viscosity behavior of concentrated therapeutic antibodies. The study used 27 FDA-approved antibodies and utilized features based on their charge, hydrophobicity, and hydrophilicity properties. In addition, Lai et al.^17^ used machine learning to predict aggregation rates of concentrated therapeutic antibodies. This study utilized 21 high-concentration therapeutic antibody with experimental aggregation rates, obtained SAP and SCM scores from MD simulations across different domains of antibodies as features and employed the feature selection method to select the best four-feature combinations. Moreover, Lai et al.^18^ used machine learning to predict antibody aggregation and viscosity at high concentrations (150 mg/ml). This study utilized 20 preclinical and clinical-stage antibodies. Despite the success of these machine-learning models, the features need to be calculated from time-consuming MD simulations.

Deep Learning is a subset of machine learning that consists of many multi-layer neural networks with many hidden units^19,20^. The common architectures include artificial neural networks (ANN), convolutional neural networks (CNN), and recurrent neural networks (RNN). Unlike traditional machine learning, deep learning can learn features by itself^21^. Deep learning has been adopted in previous studies over the years to study and predict antibody properties^22^, structures^23–25^, ability to bind to target antigen^26^, specific B-cell epitope^27^, and apparent solubility^28^. Rai et.al^29^ used deep learning to predict antibody viscosity at high concentrations using the electrostatic potential surface of the antibody variable region (Fv) as input, which still requires structural information. Lai^30^ used deep learning to develop a convolutional neural network surrogate model, DeepSCM, which requires only sequence information to predict the SCM score of antibodies in the entire Fv which can then be used to predict high concentration antibody viscosity. However, DeepSCM only accounts for the surface charges of the Fv region and its predictive capability could further be improved by including other biophysical properties across the different regions. Studies have shown that both charge (obtainable from SCM), solvent-accessible surface area, and hydrophobicity (obtainable from SAP) are key descriptors influencing the aggregation rates and viscosity of monoclonal antibodies (mAbs)^16–18^. In light of this, a promising avenue for advancing the prediction accuracy of antibody stability during early-stage drug discovery and development involves the creation of an antibody-specific sequence-based tool. Such a tool would comprehensively capture both charge and hydrophobicity, offering a more holistic approach to predicting antibody behavior.

In this study, we applied deep learning to develop DeepSP, a collection of different surrogate models that can be used to predict average dynamic SCM and SAP scores in different domains of an antibodies not just the entire variable region with a much larger and diverse datasets (N=20530) solely based on the antibody sequences, thereby accelerating MD simulations and providing a more comprehensive and holistic model for predicting antibody behavior. The sequences used for model training were obtained from the Observed Antibody Space (OAS) database^31^. First, we performed MD simulations to calculate the dynamic average and standard deviation of SAP_positive (SAP_pos), SCM_negative (SCM_neg) and SCM_positive (SCM_pos) scores in the CDRH1, CDRH2, CDRH3, CDRL1, CDRL2, CDRL3, CDR, Hv, Lv and Fv regions of these antibodies. This process yielded a total of 30 structural properties, as summarized in Table 1. We then trained a deep learning surrogate model – DeepSP, using these MD-derived averages as outputs and the preprocessed antibody sequences as inputs for model training. The relative standard deviation was utilized to quantify the error of uncertainty in the prediction of the average scores. The linear correlation coefficient of the DeepSP scores and MD-derived scores for these properties achieved values between 0.76 and 0.96 with an average of 0.87 on test set (N= 2053).

**Table 1.** List of mAb properties and domains in DeepSP model. The properties are calculated with an in-house program.

| mAb Properties | Domains |
| --- | --- |
|  | CDRH1 |
|  | CDRH2 |
|  | CDRH3 |
| Spatial aggregation propensity (SAP_pos) | CDRL1 |
| Spatial negative charge map (SCM_neg) | CDRL2 |
| Spatial positive charge map (SCM_pos) | CDRL3 |
|  | CDR |
|  | Hv |
|  | Lv |
|  | Fv |

To further validate the performance of DeepSP, we utilized a dataset comprising aggregation rates of 21 high-concentration (150 mg/mL) mAbs obtained from a previous study^17^. In this study, we employed a similar approach to the original study, using machine learning models to predict antibody aggregation rates. However, instead of using MD simulations to generate features, we utilized the DeepSP model to predict 30 structural properties of 21 antibodies, which we used as inputs (features) to train various machine learning models. We observed remarkable results, with a high correlation coefficient (R = 0.97) and low mean squared error (MSE = 0.03) between the experimental and predicted aggregation rates. Leave-one-out cross-validation (LOOCV) yielded a correlation coefficient (R = 0.75) and MSE value (MSE = 0.18). This is similar to the results obtained from the previous study that used MD simulations to generate the same features to train a machine learning model to predict their aggregation rates.

These **DeepSP** features can also serve as input in other research to train other machine learning or deep learning models to predict other desired properties of the antibodies with known and available sequences. By implementing this deep learning model during antibody screening or engineering processes, it becomes possible to identify antibodies that may have stability issues, allowing for targeted re-engineering or removal from the antibody panel.

## 2. Methods

### 2.1 Antibody sequence datasets and preprocessing

Antibody sequences were retrieved from the Observed Antibody Space – OAS^31^, Duplicated antibody sequences and those with unpaired Fv regions were removed from the dataset. The length of these antibody sequences varies and were therefore annotated with the IMGT numbering scheme using ANARCI^32^ to ensure the same input size was achieved for deep learning algorithms. The heavy chain and light chain variable region ranged from H1 to H128 and L1 to L127 respectively, with gaps filled by dashes. The maximum length allowed on the CDRH3 region [H105-H117] was 30. Sequences with insertions on other CDR or framework regions were removed from the dataset. Furthermore, sequences that do not have exactly two cysteine residues at positions 23 and 104 in the heavy and light chains were removed. Finally, sequences that failed to generate homology models for the Fv regions were excluded. This approach was adapted from a previous study^30^. These steps resulted in 23520 antibody Fv sequences.

### 2.2 Computational Modeling of mAbs and Molecular Dynamics Simulations

The homology models of the Fv regions were generated by ABodyBuilder-ML^33^. IMGT numbering was used to annotate the final models. Molecular dynamics simulations, following the procedure in previous study^30^, were conducted using all-atom antibody Fv structures in explicit solvent with the TIP3P water model^34^, VMD was used to set up a 12 Å water box beyond the protein surface^35^. CHARMM36m force field^36,37^ was implemented for MD simulations using the NAMD software package^38^. The Particle Mesh Ewald (PME) method was applied for electrostatic interactions^39^. After a gradual heating process from 100 K to 300 K, constraints were relaxed. Additional simulation details can be found in the previous study^30^. However, to confirm the appropriate equilibrium and production run time, we conducted a 10 ns equilibrium run and a 50 ns production run for the three different antibodies in our dataset to determine the optimal production run time for stabilizing the desired properties. The structural properties reached a stable state after a 10 ns production run. The simulation was then run with 10 ns equilibrium and 10 ns production runs for all antibodies, and the integration time step was set to 2 fs. Only 20530 antibody sequences made it to this stage. The final annotated CSV file, provided as Supporting Information (**annotated_oas_data.csv**), contains all included antibody sequences retrieved from the OAS database, along with reasons for excluding those that did not meet the criteria up to this stage. We proceeded to calculate the dynamic averages of SCM and SAP scores of the remaining antibodies as described in the next section.

### 2.3 Calculation of Spatial Charge Map and Spatial Aggregation Propensity Scores

The spatial charge map (SCM) is a score that was developed to differentiate low or high antibody viscosity in high concentrated solutions. The calculation of SCM scores follows previous work^1^. Briefly, the atomic SCM value has the following form.

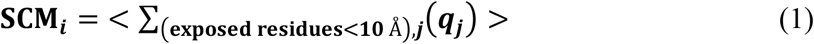

where <> indicates ensemble average from MD simulations. The atomic SCM value (*SCM*_*i*_) is the summation of all the partial charges (*qj*) on the surrounding atom *j*, which are within 10 Å of atom *i* that belongs to exposed residues. The exposed residues are considered if the sum of the side chain solvent accessible area is ≥10 Å^2^. The SCM score in different regions is then expressed as:

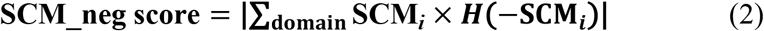

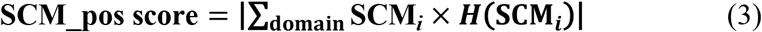

where domain refers to CDRH1, CDRH2, CDRH3, CDRL1, CDRL2, CDRL3, CDR, Hv, Lv, and Fv, ***H*** is the Heaviside function, and | . | is the absolute value function.

The spatial aggregation propensity (SAP) is a tool used to identify the location and size of aggregation-prone regions in antibodies. The calculation of SAP follows previous work^2^. The atomic SAP value is calculated as

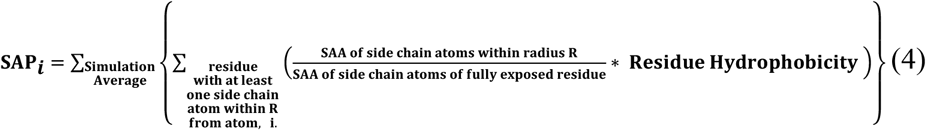

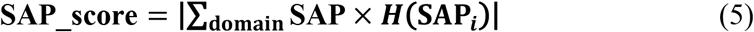

The SAP values in different regions, CDRH1, CDRH2, CDRH3, CDRL1, CDRL2, CDRL3, CDR, Hv, Lv and Fv, were also obtained.

### 2.4 Development of DeepSP using Deep Learning Models

Deep learning models were developed in Python 3.9.13 utilizing scikit-learn v1.0.2. for the train_test_split function and Keras v2.11.0 Sequential model as a wrapper for TensorFlow v2.11.0^40–42^. The CNN architecture employed in this study consisted of three convolutional layers, each integrated with batch normalization and dropout layers, followed by a pooling layer, flattening operation, and a densely connected layer with a single output layer.

Hyperparameter optimization was performed using the Keras Tuner library^43^ with three different optimization techniques - Hyperband, Random Search, and Bayesian Optimization techniques^44^ to efficiently explore the hyperparameter space and identify the best-performing configurations for the neural network model. Various combinations of hyperparameters were explored, including the number of filters (ranging from 16 to 128 with increments of 16), kernel sizes (selected from [3, 4, 5]), dropout rates (ranging from 0.0 to 0.5 in steps of 0.1), number of units in dense layers (ranging from 32 to 128 with a step of 16), and learning rates (chosen from [1e-2, 5e-3, 1e-3, 1e-4]). The optimal configuration was determined based on the MAE values of the best validation model.

The dataset for regression was divided into training (65%), validation (25%), and test sets (10%). The best hyperparameters were obtained from keras tuner over 50 epochs with a batch size of 32 and the Adam optimizer. The best model, which is the model with the minimum validation loss was saved using Model Checkpoint from keras.callbacks, and the CNN architecture and weights were saved in JSON and HDF5 formats, respectively. The activation function used for the CNN model was ReLU.

In our study, two different approaches were employed to predict the spatial properties in antibodies using the methods described above. First, we trained individual CNN models for each of the 30 properties, resulting in a total of 30 models in our DeepSP collection. Second, we trained three models, each predicting a property (SAP_pos, SCM_pos, or SCM_neg) across all 10 regions of the antibodies, resulting in 10 outputs per model. For instance, the SAP_pos model can predict properties across different antibody regions, such as SAP_pos_CDRH1, SAP_pos_CDRH2, and so on. No significant differences were observed in the predictions after comparing the outcomes of these two approaches. Detailed information on best and standard deviation scores hyperparameters and model performance metrics for both approaches can be found in Tables S1-S4 in the Supporting Information. We decided to adopt the latter approach for the rest of the project.

### 2.5 Machine learning feature selection and modeling to predict aggregation rate using DeepSP features

To validate the performance of the DeepSP model established in this study, we utilized a dataset comprising aggregation rates of 21 high-concentration (150 mg/mL) mAbs obtained from previous research^17^. DeepSP was used to generate 30 structural properties as features in machine learning model training. Given the limited dataset size, the risk of overfitting^45^ arises when dealing with numerous features. Therefore, we applied the Exhaustive Feature Selector (EFS) tool from mlxtend library^46^ in conjunction with various regression algorithms for feature selection. We iteratively assessed different feature subsets based on the negative mean squared error as the scoring metric, varying feature counts, and cross-validation folds. Subsequently, we computed the mean MSE for specific feature subsets identified by the EFS. For each subset, MSE was computed using different regression models using a repeated k-fold cross-validation method.

Finally, we collected all subset details and their associated averaged MSE values, selecting the feature combination with the smallest MSE value to train the machine learning model.

The machine learning algorithms from the scikit-learn library^40^ used are linear regression (LR), k-nearest neighbors regressor (KNN), support vector regressor (SVR) and random forest regressor (RFR). After selecting the best feature combinations obtained from the exhaustive feature combination, each machine learning model was trained and tuned to obtain the optimal hyperparameter that will give the best model. For KNN, we varied the number of neighbors from 2 to 8, for SVR, we tuned the parameters C (ranging from 5.0 to 15.0) and ε (ranging from 0.1 to 0.5), while for RFR, we adjusted the max_depth parameter (ranging from 2 to 6). We then evaluated each model’s performance by comparing the correlation coefficients (r) and MSE between the experimental and predicted data. The model that exhibited the highest correlation coefficients and the lowest MSE was selected as our final machine-learning model. To verify the reliability of our models, we implemented LOOCV, a commonly used technique in machine learning and statistics for model performance assessment, particularly in situations with limited data. While tuning the parameters, we concurrently created a validation model using LOOCV with the same set of parameters. Although the correlation coefficients and mean square error of the validation model exhibited slight reductions compared to the initial model, we established a threshold. If the correlation coefficients did not decrease by more than 0.3, we considered the model as yielding reliable results.

## 3. Results and Discussions

### 3.1 Antibody sequence dataset and statistical analysis

The antibody variable region paired sequences (30,000) were retrieved from OAS^31^. The preprocessing steps (detailed in the Materials and Methods section), which includes filtering out sequences based on some criteria such as complementarity determining region (CDR) length, the number of cysteine residues, and insertion yielded 25320 sequences and after removing the unstable ones during MD simulations, 20530 antibody Fv sequences were left for this study.

Figure 2 shows the length distribution of different antibody regions in the dataset. The VH length and VL length were approximately normally distributed, centered at 122 and 108 respectively. The first complementarity determining region of the heavy chain (CDRH1) length had the highest peak at 7 which constitute about 85% of the data set, and the rest had length of 8-9. The first complementarity determining region of the light chain (CDRL1) length had the highest peak at 7 which constitutes about 70% of the dataset and the rest had length of 9. For the second complementarity determining region of the heavy chain (CDRH2) length, the highest peak was at 6, and the second-highest peak was at 5 and the rest has 7 or 8. The second complementary determining region of the light chain (CDRL2) length had the highest peak at 6 and the rest has length of 5. For the third complementarity determining region of the light chain (CDRL3) length, the highest peak was at 15. The third complementary determining region of the heavy chain (CDRH3) length had a wide distribution centered at 13.

**Figure 1.**
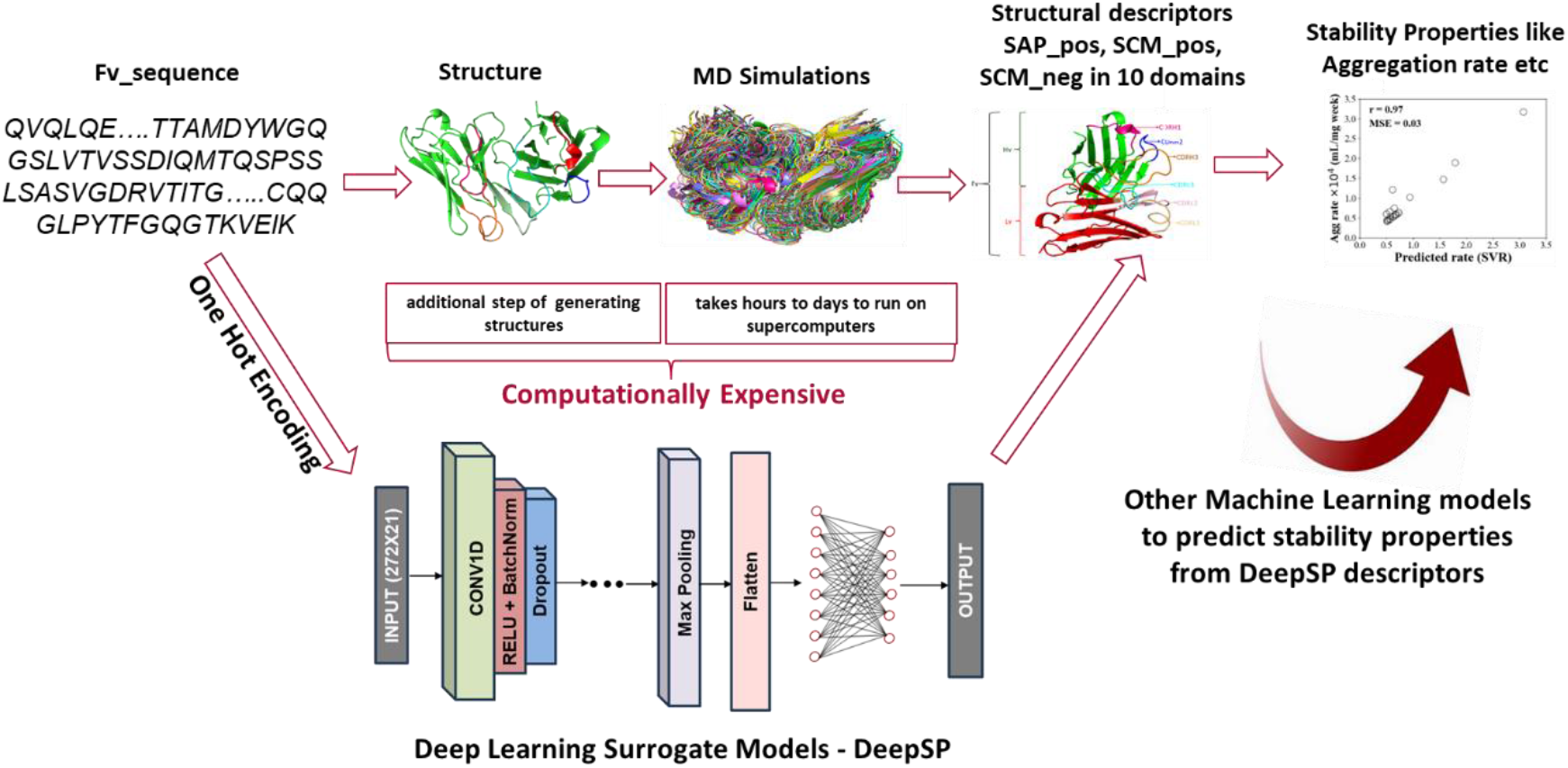
Illustration of the workflow to accelerate MD simulations using deep learning and consequently predicting the stability at high concentration.

**Figure 2.**
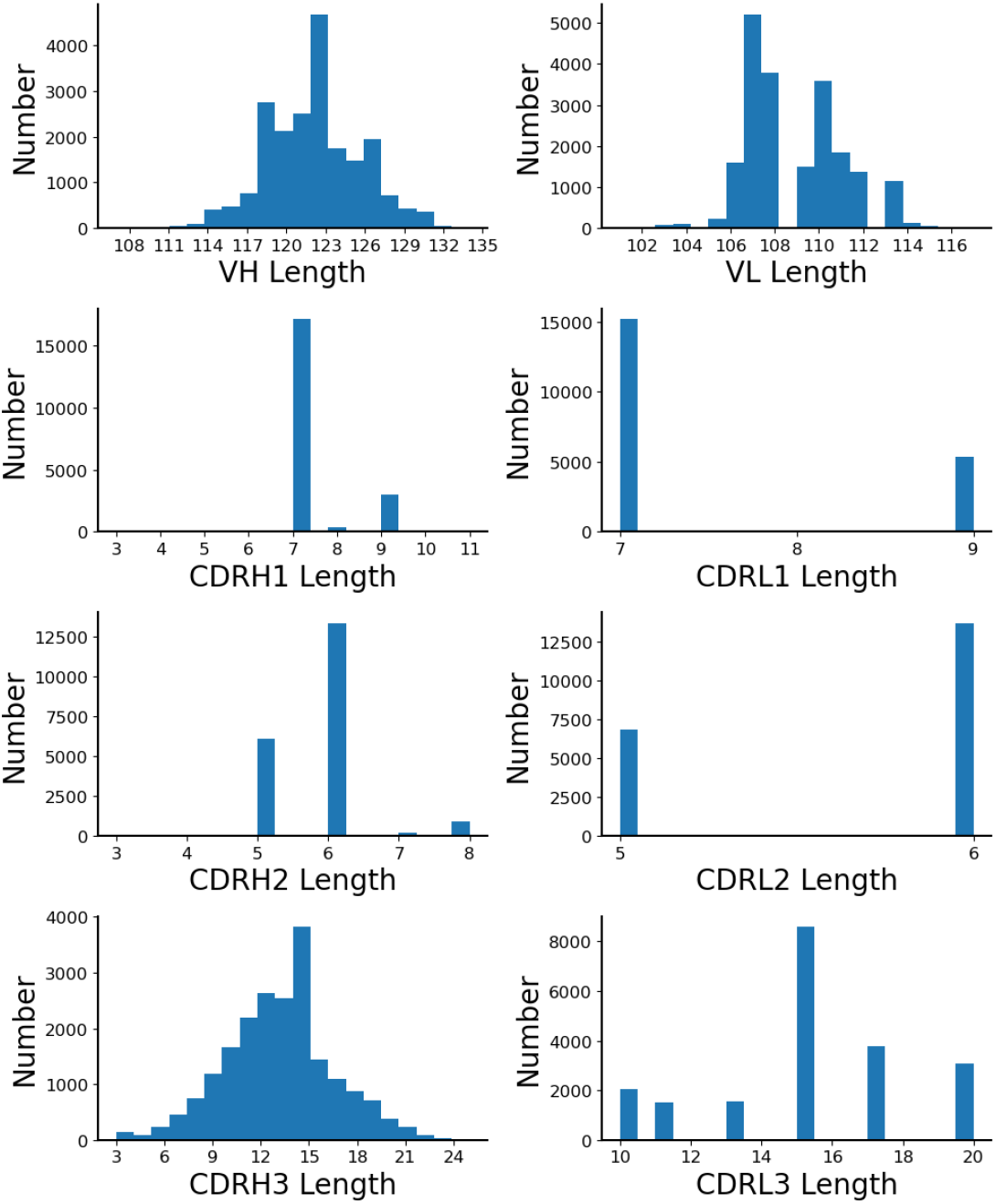
Distribution of VH, VL, CDRH1, CDRH2, CDRH3, CDRL1, CDRL2, and CDRL3 lengths of the 20530 Fv sequences in this study. The CDR regions are based on the Chothia definition.

CNN models require the input to have a fixed size, however, our antibody sequences have variable lengths. To address the variable length issue, we adopted the Chothia numbering scheme^47^ to annotate the heavy and light chain variable regions. This choice was made over IMGT due to Chothia’s focus on conserved CDRs, enabling better alignment and representation of antibody functionality^48^. With Chothia, we ensured the CDRs were accurately captured, allowing for precise analysis and modeling of antibody properties. Gaps were padded with dashes, resulting in fixed lengths of 145 and 127 for the heavy and light chain variable regions, respectively.

### 3.2 MD simulations, SCM and SAP calculation of the antibody in the dataset

The homology models of the 20530 antibody variable regions were constructed and prepared to perform MD simulations. The SAP_pos, SCM_pos and SCM_neg scores were calculated in the CDRH1, CDRH2, CDRH3, CDRL1, CDRL2, CDRL3, CDR, Hv, Lv, and Fv regions by the ensemble averages over 10 ns. Figure 3 shows the box-and- whisker of the normalized average and standard deviation score distribution for all the 30 spatial properties as obtained from MD simulation. Table S5 summarizes the analysis of the 30 spatial properties.

**Figure 3.**
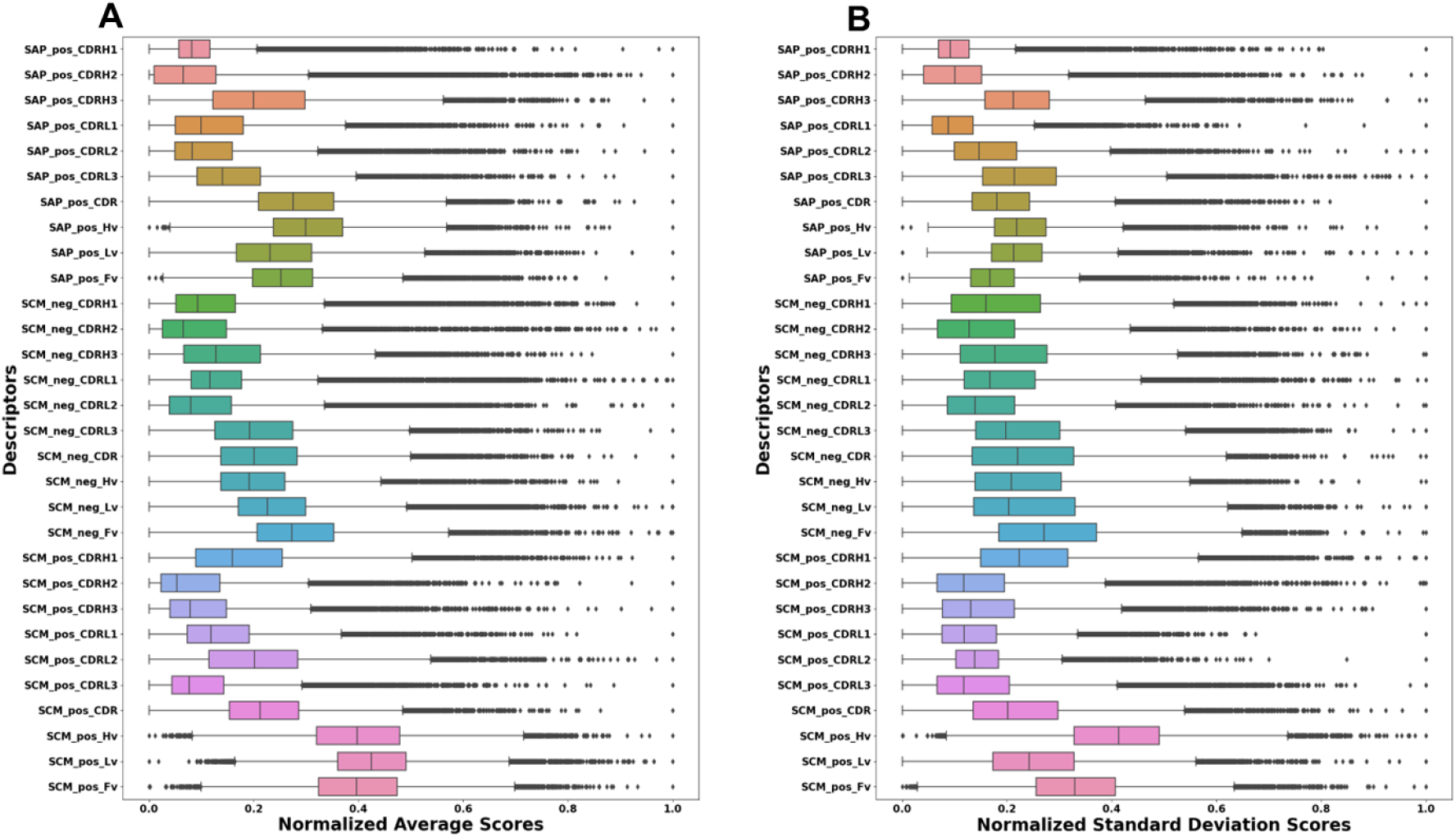
Box-and-Whisker plot illustrating the normalized A) average B) standard deviation score distribution for all 30 properties obtained from MD simulations.

The simulation run was initially conducted for 10 ns equilibrium and 50 ns production run to test the convergence. Figure S1 shows the time trajectory plot of the SAP_pos, SCM_pos, SCM_neg scores in the Hv and Fv regions of an antibody. The scores fluctuated around the mean, and the mean converged and stabilized after 10 ns production run, hence 10 ns equilibrium run, and 10 ns production was maintained for the other antibodies. Unlike in full-length antibodies, which demand extended simulation times for convergence and stability, single-variable region simulations achieve quicker equilibration which makes them more suitable for high-throughput computing in large antibody datasets.

### 3.3 CNN model training and optimization for the DeepSP model

CNN model was chosen for model development in this study as it has been shown to perform better than other deep learning models like ANN and RNN for predicting antibody binders [17]. The ratio for training/validation/test split was 65:25:10. The architecture and parameters were optimized by tuning hyperparameters using keras tuner (as detailed in the Materials and Methods section). The MAE, validation loss, and correlation between the prediction and actual scores for the 10% test set were used to evaluate and select the best-performing model.

Figure 4 shows the CNN architecture of the three models. Each model had an input shape of (272, 21). The number of columns is the sum of heavy chain variable region length (145) and light chain variable region length (127). The rows came from one-hot encoding, including 20 amino acids and one gap. The input layer was connected to a 1D CNN layer using the activation function of the rectified linear unit (Relu). Following the architecture illustrated in the figure, the fully connected layer was connected to an output layer of size 10. Figure 4 shows the training and validation loss over the training epochs for all three models. While the training and validation loss generally converge, there is noticeable divergence in the case of SCM_pos after 20 epochs. However, we used model checkpoint from keras callback to monitor the model performance and implement the one with the best hyperparameter.

**Figure 4.**
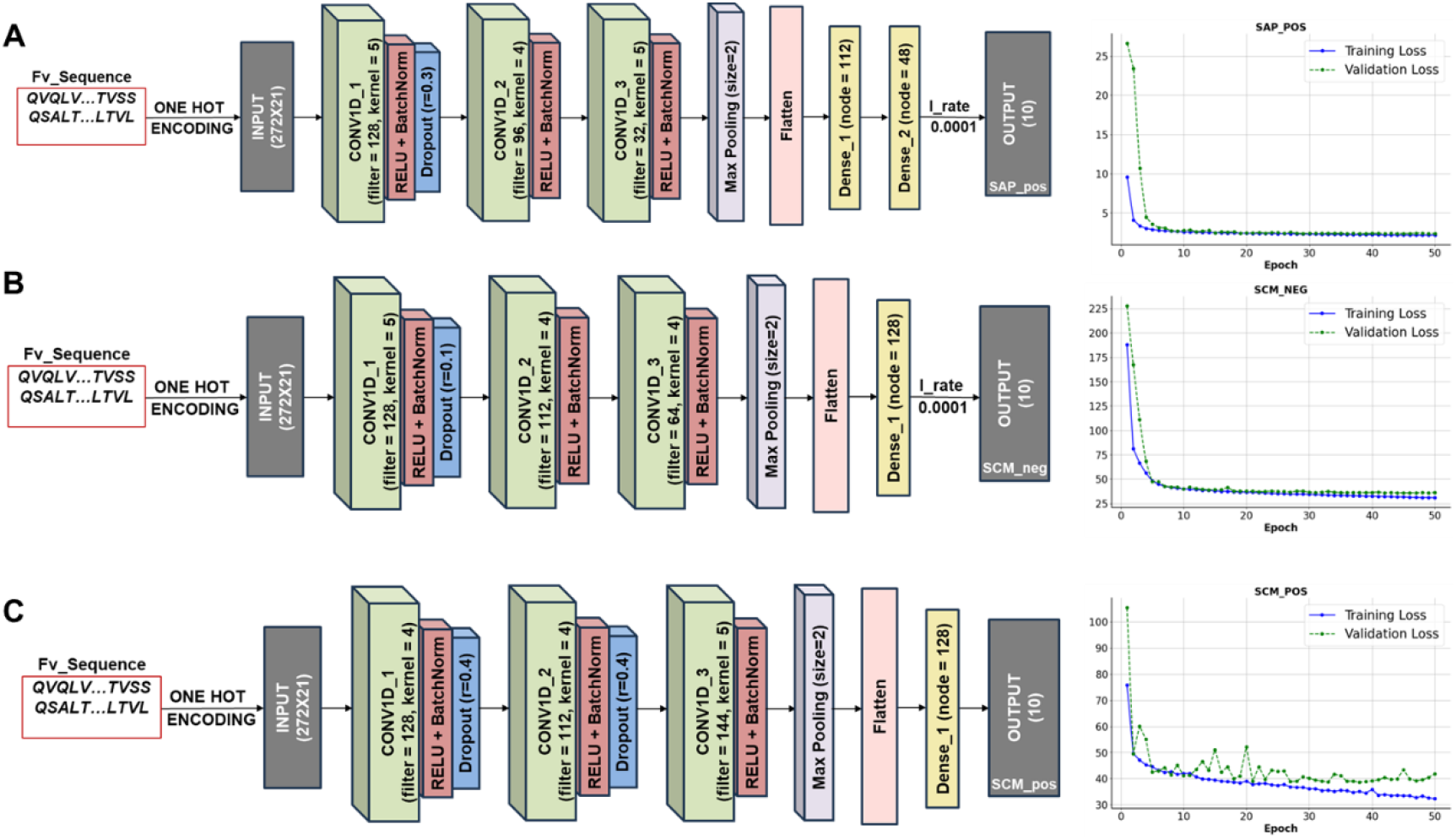
Illustration of CNN model architecture with the training and validation loss over number of epochs for A) SAP_pos model B) SCM_neg model C) SCM_pos model, contained in DeepSP surrogate model developed in this study.

Table S3 shows the best and optimal hyperparameter combination generated from keras tuner. For each model, the model with the minimum MAE, minimum validation loss and maximum correlation between actual and predicted test set was selected. Table S4 shows the model performance metrics for the final models. Figure 5 shows the correlation between the predicted and actual scores for all properties. A minimum correlation of 76% and maximum correlation of 97%, and the MAE of all the properties greatly beats the baseline MAE as shown in scatter plot illustrated in Figure S2. The relative standard deviation was obtained by dividing the standard deviation by the actual value and can be found for each property in Figure S3.

**Figure 5.**
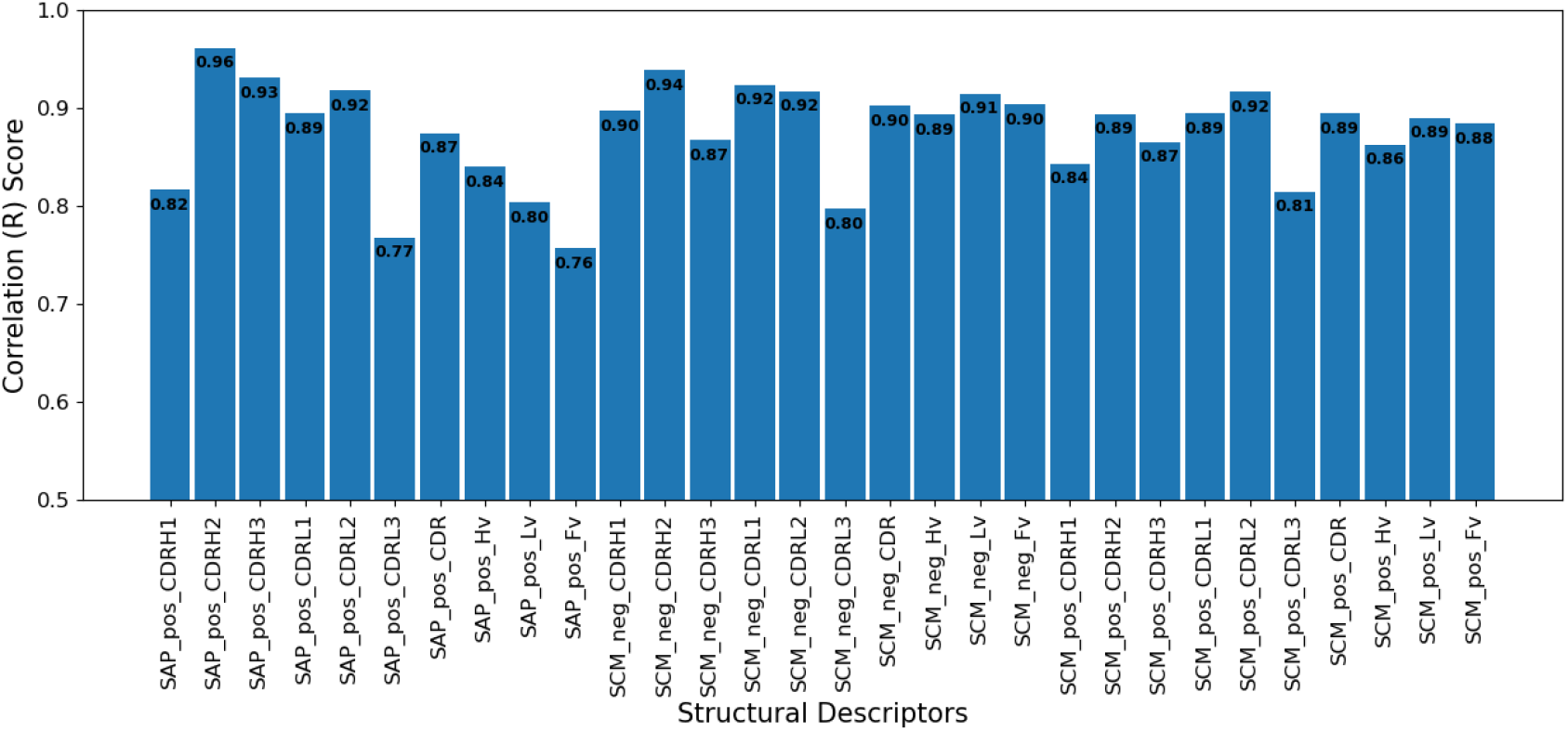
Bar plot illustrating the correlation between the predicted and actual score of all 30 spatial properties.

### 3.4 Aggregation Rate Prediction using DeepSP model as features

In the previous study^17^, a machine-learning model was proposed to predict the aggregation rates of 21 mAbs. MD simulation was employed to compute the SAP and SCM in different mAb domains, which served as features for machine learning model training. Here, we used DeepSP developed in this study to predict these features solely from the Fv sequences, as demonstrated in Table 1. This is to validate the ability of DeepSP features to be able to alleviate the computationally expensive MD simulation in generating these features for training machine learning models to predict the aggregation rate of an antibody.

For feature selection, various feature combinations were explored for Linear Regression, Support Vector Regression (SVR), Random Forest (RF), and K-Nearest Neighbors (KNN) models using 4-fold cross-validation. Subsequently, we computed the MSE values for machine learning models built using different feature combinations. These MSE values were used to determine the optimal set of features for subsequent machine learning model training.

In Table 2, we listed the MSE values calculated for three and four feature combinations in order to determine the feature combinations with the lowest MSE values to be used, followed by hyperparameter tuning for the models. This approach ensures that we can effectively select the best feature combinations when training machine learning models, thereby improving the predictive and generalization capabilities of the models.

**Table 2.** Mean squared error (MSE) of the best three-feature and four-feature combinations of the linear regression, support vector regression (SVR), k-nearest neighbors’ regression (KNN), and random forest regression (RF) models for predicting aggregation rates.

| Regression Models | Three-feature | MSE | Four-feature | MSE |
| --- | --- | --- | --- | --- |
| Linear | SAP_pos_CDRH3 | 0.433 | SAP_pos_CDRH3 | 0.457 |
|  | SCM_pos_CDRL3 |  | SCM_pos_CDRL1 |  |
|  | SCM_neg_CDRH3 |  | SCM_pos_CDR |  |
|  |  |  | SCM_pos_Hv |  |
| Nearest neighbors | SAP_pos_CDRL3 | 0.366 | SCM_pos_CDRH3 | 0.319 |
|  | SCM_pos_CDRH3 |  | SCM_neg_CDRH2 |  |
|  | SCM_neg_Fv |  | SCM_neg_Hv |  |
|  |  |  | SCM_neg_Fv |  |
| Random forest | SAP_pos_Hv | 0.367 | SAP_pos_Hv | 0.364 |
|  | SCM_pos_CDRH3 |  | SCM_pos_CDRH3 |  |
|  | SCM_neg_Fv |  | SCM_pos_Lv |  |
|  |  |  | SCM_neg_Fv |  |
| Support vector | SCM_pos_CDRH3 | 0.307 | SCM_pos_CDRH3 | 0.301 |
|  | SCM_neg_CDRH2 |  | SCM_neg_CDRH2 |  |
|  | SCM_neg_Fv |  | SCM_neg_CDRL2 |  |
|  |  |  | SCM_neg_Fv |  |

After tuning the hyperparameters, Table 3 summarizes the results of the best three-feature or four-feature combinations of Linear Regression, SVR, RF, and KNN models. The SVR model has the highest correlation coefficient of 0.97 and a MSE of 0.03 when comparing the experimental data to the predicted data. (For comparisons with other regression models, refer to Figure S4.) To validate the reliability of our training results, we employed LOOCV on our limited dataset. LOOCV offers the advantage of maximizing the utility of available data for both training and validation, rendering it an unbiased estimator of a model’s performance. LOOCV provides a dependable estimate of a model’s performance and is particularly valuable for detecting issues like overfitting, especially in scenarios with small datasets where leveraging available information is crucial. Table 3 summarizes the performance of all regression models in comparison to their LOOCV performance which shows that SVR model, evaluated using LOOCV, yielded a correlation coefficient (r) of 0.75 and a MSE of 0.18. While the correlation coefficient exhibited a slight reduction, it remained within acceptable limits. Furthermore, when compared to other regression models, the SVR model outperformed them.

**Table 3.** Performance metrics, correlation coefficients (r) and mean square error (MSE) of different regression models.

| Regression Models | Features | r (all) | r (LOOCV) | MSE (all) | MSE (LOOCV) |
| --- | --- | --- | --- | --- | --- |
| Linear | SAP_pos_CDRH3 | 0.49 | 0.14 | 0.31 | 0.40 |
|  | SAP_pos_CDRL3 |  |  |  |  |
|  | SCM_neg_CDRH3 |  |  |  |  |
| Nearest neighbors<br>(neighbor numbers<br>= 3) | SCM_pos_CDRH3 | 0.85 | 0.64 | 0.11 | 0.24 |
|  | SCM_neg_CDRH2 |  |  |  |  |
|  | SCM_neg_Hv |  |  |  |  |
|  | SCM_neg_Fv |  |  |  |  |
| Random forest<br>(max_depth = 6) | SAP_pos_Hv | 0.94 | 0.47 | 0.05 | 0.32 |
|  | SCM_pos_CDRH3 |  |  |  |  |
|  | SCM_pos_Lv |  |  |  |  |
|  | SCM_neg_Fv |  |  |  |  |
| Support vector<br>(C = 15.0, $\epsilon$ = 0.1) | SCM_pos_CDRH3 | 0.97 | 0.75 | 0.03 | 0.18 |
|  | SCM_neg_CDRH2 |  |  |  |  |
|  | SCM_neg_Fv |  |  |  |  |

These results closely align with the performance obtained in the previous study^17^ where features were derived from MD simulations as shown in Figure 6. This demonstrates the effectiveness of our newly established DeepSP model, which can effectively replace the MD-based methods.

**Figure 6.**
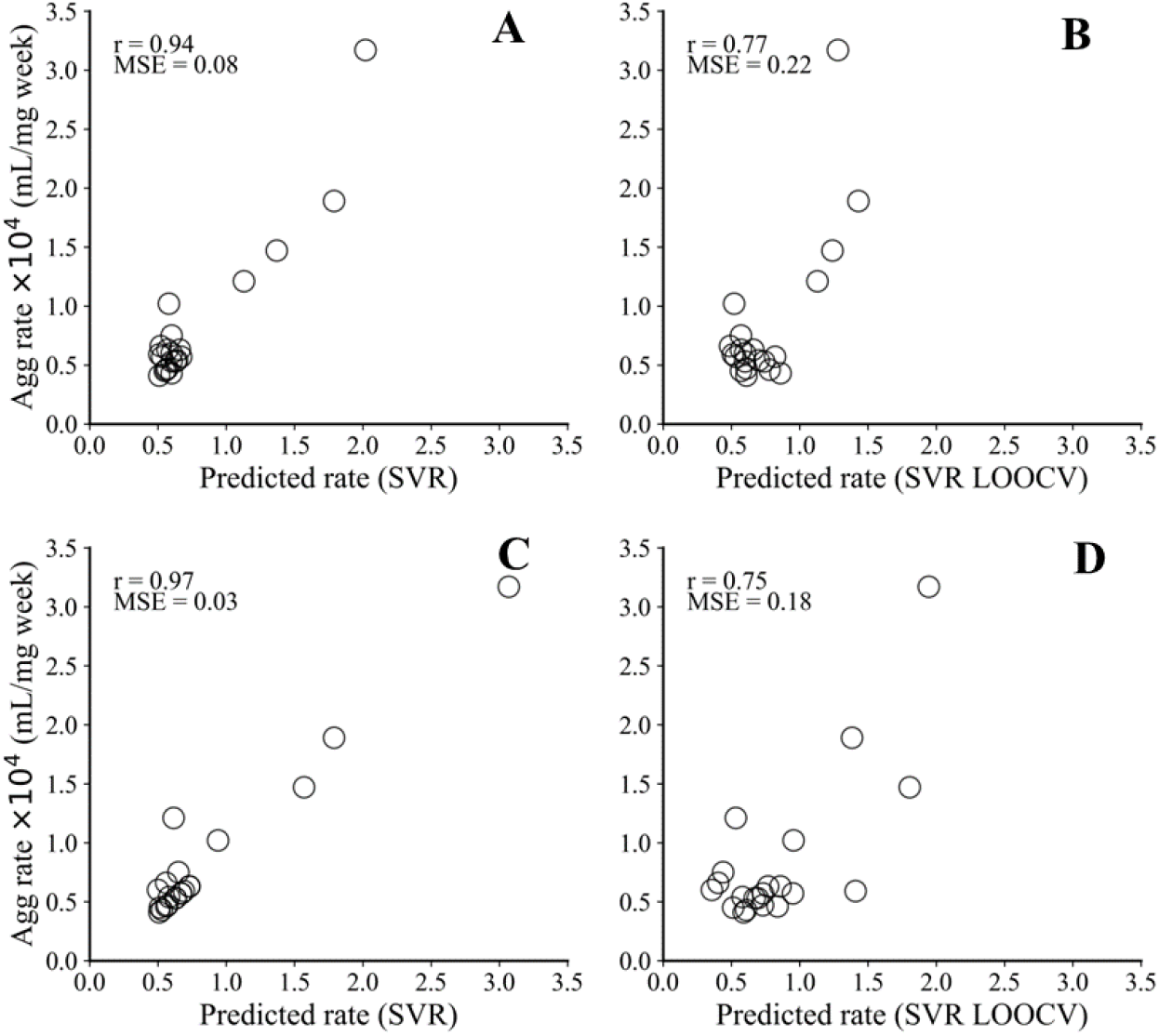
Scatter plot of correlation between predicted and experimental aggregation rate A, B) from previous study where MD simulation features were used C, D) current study where DeepSP features were used.

### 3.5 Availability and implementation of the DeepSP model

DeepSP’s source code and pretrained parameters are freely available at https://github.com/Lailabcode/DeepSP. DeepSP can be utilized using the notebook file – DeepSP_predictor.ipynb. This notebook file requires only one input which is a csv file that contains the names, heavy chain, and light chain (Fv sequences only) of the antibody whose descriptors are to be generated (see DeepSP_input.csv for sample format). Upon running this notebook file, the input antibody sequence (combined heavy and light chain) will be aligned (using ANARCI), preprocessed, and encoded. The DeepSP saved model in JSON and h5 format will then be used to generate the predicted scores of all the 30 spatial properties, displayed as a dataframe and saved to a csv file. The python file - DeepSP_train.py contains the code that was used for DeepSP training, validation, and testing.

## 4. Conclusion

DeepSP was developed as a surrogate model to accelerate the MD simulation-based tools for calculating SAP and SCM scores in all 6 CDR regions, the entire CDR region, Hv, Lv, and the entire Fv region of an antibody solely from the sequence. It was trained using high-throughput MD simulation results and 1D convolutional neural network architecture. DeepSP, as an antibody-specific model, incorporates features such as charge and hydrophobicity. This makes it a more comprehensive descriptor for antibodies, enhancing its capability to predict and assess antibody stability accurately. DeepSP has been used to predict spatial properties, which served as input or features to an SVR model, trained to predict the aggregation rate of 21 monoclonal antibodies.

DeepSP features can serve as antibody-specific features for training machine learning models for other stability properties such as viscosity (manuscript in preparation) and solubility as well as other desired properties using only Fv sequences. These tools can screen for hundreds of antibody drug candidates within a few seconds. The DeepSP features can also be used to train surrogate models for other biophysical properties from experiments such as melting temperature, retention time from hydrophobic interaction chromatography, etc. Deep learning paves a promising way for predicting antibody functions to facilitate drug design. Overall, this tool will facilitate early-stage drug development.

## Abbreviations

CDR: complementarity determining region
CDRH1: the first complementarity determining region of the heavy chain
CDRH2: the second complementarity determining region of the heavy chain
CDRH3: the third complementarity determining region of the heavy chain
CDRL1: the first complementarity determining region of the light chain
CDRL2: the second complementarity determining region of the light chain
CDRL3: the third complementarity determining region of the light chain
Fv: variable fragment
mAbs: monoclonal antibodies
MD: molecular dynamics
SCM: spatial charge map
SAP: spatial aggregation propensity
VH: heavy chain variable region
VL: light chain variable region

## Acknowledgments

This work was financially supported by start-up funds from Stevens Institute of Technology. We thank the Advanced Cyberinfrastructure Coordination Ecosystem: Services & Support (CHM210016) and the Oak Ridge Leadership Computing Facility (BIP232) for supporting computing resources.

## Notes

https://github.com/Lailabcode/DeepSP

### Competing Interest Statement

The authors have declared no competing interest.

## Reference

(1) Agrawal, N. J.; Helk, B.; Kumar, S.; Mody, N.; Sathish, H. A.; Samra, H. S.; Buck, P. M.; Li, L.; Trout, B. L. Computational Tool for the Early Screening of Monoclonal Antibodies for Their Viscosities. MAbs 2016, 8 (1), 43–48. 10.1080/19420862.2015.1099773.

(2) Chennamsetty, N.; Voynov, V.; Kayser, V.; Helk, B.; Trout, B. L. Design of Therapeutic Proteins with Enhanced Stability. Proceedings of the National Academy of Sciences 2009, 106 (29), 11937–11942. 10.1073/pnas.0904191106.

(3) Bhambhani, A.; Kissmann, J. M.; Joshi, S. B.; Volkin, D. B.; Kashi, R. S.; Middaugh, C. R. Formulation Design and High-Throughput Excipient Selection Based on Structural Integrity and Conformational Stability of Dilute and Highly Concentrated IgG1 Monoclonal Antibody Solutions. J Pharm Sci 2012, 101 (3), 1120–1135. 10.1002/jps.23008.

(4) Shire, S. J.; Shahrokh, Z.; Liu, J. Challenges in the Development of High Protein Concentration Formulations. J Pharm Sci 2004, 93 (6), 1390–1402. 10.1002/jps.20079.

(5) Berteau, C.; Filipe-Santos, O.; Wang, T.; Rojas, H. E.; Granger, C.; Schwarzenbach, F. Evaluation of the Impact of Viscosity, Injection Volume, and Injection Flow Rate on Subcutaneous Injection Tolerance. Med Devices (Auckl) 2015, 8, 473–484. 10.2147/MDER.S91019.

(6) Zhang, Z.; Liu, Y. Recent Progresses of Understanding the Viscosity of Concentrated Protein Solutions. Current Opinion in Chemical Engineering 2017, 16, 48–55. 10.1016/j.coche.2017.04.001.

(7) Viola, M.; Sequeira, J.; Seiça, R.; Veiga, F.; Serra, J.; Santos, A. C.; Ribeiro, A. J. Subcutaneous Delivery of Monoclonal Antibodies: How Do We Get There? J Control Release 2018, 286, 301–314. 10.1016/j.jconrel.2018.08.001.

(8) Matucci, A.; Vultaggio, A.; Danesi, R. The Use of Intravenous versus Subcutaneous Monoclonal Antibodies in the Treatment of Severe Asthma: A Review. Respir Res 2018, 19, 154. 10.1186/s12931-018-0859-z.

(9) Jiskoot, W.; Hawe, A.; Menzen, T.; Volkin, D. B.; Crommelin, D. J. A. Ongoing Challenges to Develop High Concentration Monoclonal Antibody-Based Formulations for Subcutaneous Administration: Quo Vadis? J Pharm Sci 2022, 111 (4), 861–867. 10.1016/j.xphs.2021.11.008.

(10) Xie, M. How Do You Obtain the Sequence of an Antibody?. Rapid Novor. https://www.rapidnovor.com/how-obtain-sequence-antibody/ (accessed 2024-02-20).

(11) Chaudhri, A.; Zarraga, I. E.; Kamerzell, T. J.; Brandt, J. P.; Patapoff, T. W.; Shire, S. J.; Voth, G. A. Coarse-Grained Modeling of the Self-Association of Therapeutic Monoclonal Antibodies. J Phys Chem B 2012, 116 (28), 8045–8057. 10.1021/jp301140u.

(12) Chowdhury, A.; Bollinger, J. A.; Dear, B. J.; Cheung, J. K.; Johnston, K. P.; Truskett, T. M. Coarse-Grained Molecular Dynamics Simulations for Understanding the Impact of Short-Range Anisotropic Attractions on Structure and Viscosity of Concentrated Monoclonal Antibody Solutions. Mol Pharm 2020, 17 (5), 1748–1756. 10.1021/acs.molpharmaceut.9b00960.

(13) Izadi, S.; Patapoff, T. W.; Walters, B. T. Multiscale Coarse-Grained Approach to Investigate Self-Association of Antibodies. Biophys J 2020, 118 (11), 2741–2754. 10.1016/j.bpj.2020.04.022.

(14) Lai, P.-K.; Swan, J. W.; Trout, B. L. Calculation of Therapeutic Antibody Viscosity with Coarse-Grained Models, Hydrodynamic Calculations and Machine Learning-Based Parameters. mAbs 2021, 13 (1), 1907882. 10.1080/19420862.2021.1907882.

(15) Wang, G.; Varga, Z.; Hofmann, J.; Zarraga, I. E.; Swan, J. W. Structure and Relaxation in Solutions of Monoclonal Antibodies. J Phys Chem B 2018, 122 (11), 2867–2880. 10.1021/acs.jpcb.7b11053.

(16) Lai, P.-K.; Fernando, A.; Cloutier, T. K.; Gokarn, Y.; Zhang, J.; Schwenger, W.; Chari, R.; Calero-Rubio, C.; Trout, B. L. Machine Learning Applied to Determine the Molecular Descriptors Responsible for the Viscosity Behavior of Concentrated Therapeutic Antibodies. Mol Pharm 2021, 18 (3), 1167–1175. 10.1021/acs.molpharmaceut.0c01073.

(17) Lai, P.-K.; Fernando, A.; Cloutier, T. K.; Kingsbury, J. S.; Gokarn, Y.; Halloran, K. T.; Calero-Rubio, C.; Trout, B. L. Machine Learning Feature Selection for Predicting High Concentration Therapeutic Antibody Aggregation. J Pharm Sci 2021, 110 (4), 1583–1591. 10.1016/j.xphs.2020.12.014.

(18) Lai, P.-K.; Gallegos, A.; Mody, N.; Sathish, H. A.; Trout, B. L. Machine Learning Prediction of Antibody Aggregation and Viscosity for High Concentration Formulation Development of Protein Therapeutics. MAbs 2022, 14 (1), 2026208. 10.1080/19420862.2022.2026208.

(19) Sarker, I. H. Deep Learning: A Comprehensive Overview on Techniques, Taxonomy, Applications and Research Directions. SN Comput Sci 2021, 2 (6), 420. 10.1007/s42979-021-00815-1.

(20) Emmert-Streib, F.; Yang, Z.; Feng, H.; Tripathi, S.; Dehmer, M. An Introductory Review of Deep Learning for Prediction Models With Big Data. Frontiers in Artificial Intelligence 2020, 3.

(21) Kansara, D., Sawant, V. (2020). Comparison of Traditional Machine Learning and Deep Learning Approaches for Sentiment Analysis. In: Vasudevan, H., Michalas, A., Shekokar, N., Narvekar, M. (eds) Advanced Computing Technologies and Applications. Algorithms for Intelligent Systems. Springer, Singapore. 10.1007/978-981-15-3242-9_35 -Google Search. https://www.google.com/search?q=Kansara%2C+D.%2C+Sawant%2C+V.+%282020%29.+Comparison+of+Traditional+Machine+Learning+and+Deep+Learning+Approaches+for+Sentiment+Analysis.+In%3A+Vasudevan%2C+H.%2C+Michalas%2C+A.%2C+Shekokar%2C+N.%2C+Narvekar%2C+M.+%28eds%29+Advanced+Computing+Technologies+and+Applications.+Algorithms+for+Intelligent+Systems.+Springer%2C+Singapore.+https%3A%2F%2Fdoi.org%2F10.1007%2F978-981-15-3242-9_35&sca_esv=c5fb69bdefab0649&rlz=1C1GCEB_enUS1031US1031&ei=FdXUZfTcJdLtptQPv7ONsAM&ved=0ahUKEwi0u6SDrbqEAxXStokEHb9ZAzYQ4dUDCBA&uact=5&oq=Kansara%2C+D.%2C+Sawant%2C+V.+%282020%29.+Comparison+of+Traditional+Machine+Learning+and+Deep+Learning+Approaches+for+Sentiment+Analysis.+In%3A+Vasudevan%2C+H.%2C+Michalas%2C+A.%2C+Shekokar%2C+N.%2C+Narvekar%2C+M.+%28eds%29+Advanced+Computing+Technologies+and+Applications.+Algorithms+for+Intelligent+Systems.+Springer%2C+Singapore.+https%3A%2F%2Fdoi.org%2F10.1007%2F978-981-15-3242-9_35&gs_lp=Egxnd3Mtd2l6LXNlcnAi2QJLYW5zYXJhLCBELiwgU2F3YW50LCBWLiAoMjAyMCkuIENvbXBhcmlzb24gb2YgVHJhZGl0aW9uYWwgTWFjaGluZSBMZWFybmluZyBhbmQgRGVlcCBMZWFybmluZyBBcHByb2FjaGVzIGZvciBTZW50aW1lbnQgQW5hbHlzaXMuIEluOiBWYXN1ZGV2YW4sIEguLCBNaWNoYWxhcywgQS4sIFNoZWtva2FyLCBOLiwgTmFydmVrYXIsIE0uIChlZHMpIEFkdmFuY2VkIENvbXB1dGluZyBUZWNobm9sb2dpZXMgYW5kIEFwcGxpY2F0aW9ucy4gQWxnb3JpdGhtcyBmb3IgSW50ZWxsaWdlbnQgU3lzdGVtcy4gU3ByaW5nZXIsIFNpbmdhcG9yZS4gaHR0cHM6Ly9kb2kub3JnLzEwLjEwMDcvOTc4LTk4MS0xNS0zMjQyLTlfMzVIAFAAWABwAHgAkAEAmAEAoAEAqgEAuAEDyAEA-AEB&sclient=gws-wiz-serp (accessed 2024-02-20).

(22) Graves, J.; Byerly, J.; Priego, E.; Makkapati, N.; Parish, S. V.; Medellin, B.; Berrondo, M. A Review of Deep Learning Methods for Antibodies. Antibodies (Basel) 2020, 9 (2), 12. 10.3390/antib9020012.

(23) Ruffolo, J. A.; Guerra, C.; Mahajan, S. P.; Sulam, J.; Gray, J. J. Geometric Potentials from Deep Learning Improve Prediction of CDR H3 Loop Structures. Bioinformatics 2020, 36 (Suppl 1), i268–i275. 10.1093/bioinformatics/btaa457.

(24) Ruffolo, J. A.; Sulam, J.; Gray, J. J. Antibody Structure Prediction Using Interpretable Deep Learning. Patterns 2022, 3 (2), 100406. 10.1016/j.patter.2021.100406.

(25) Ruffolo, J. A.; Chu, L.-S.; Mahajan, S. P.; Gray, J. J. Fast, Accurate Antibody Structure Prediction from Deep Learning on Massive Set of Natural Antibodies. Nat Commun 2023, 14 (1), 2389. 10.1038/s41467-023-38063-x.

(26) Mason, D. M.; Friedensohn, S.; Weber, C. R.; Jordi, C.; Wagner, B.; Meng, S. M.; Ehling, R. A.; Bonati, L.; Dahinden, J.; Gainza, P.; Correia, B. E.; Reddy, S. T. Optimization of Therapeutic Antibodies by Predicting Antigen Specificity from Antibody Sequence via Deep Learning. Nat Biomed Eng 2021, 5 (6), 600–612. 10.1038/s41551-021-00699-9.

(27) Sher, G.; Zhi, D.; Zhang, S. DRREP: Deep Ridge Regressed Epitope Predictor. BMC Genomics 2017, 18 (6), 676. 10.1186/s12864-017-4024-8.

(28) Feng, J.; Jiang, M.; Shih, J.; Chai, Q. solPredict: Antibody Apparent Solubility Prediction from Sequence by Transfer Learning. bioRxiv December 9, 2021, p 2021.12.07.471655. 10.1101/2021.12.07.471655.

(29) Rai, B. K.; Apgar, J. R.; Bennett, E. M. Low-Data Interpretable Deep Learning Prediction of Antibody Viscosity Using a Biophysically Meaningful Representation. Sci Rep 2023, 13 (1), 2917. 10.1038/s41598-023-28841-4.

(30) Lai, P.-K. DeepSCM: An Efficient Convolutional Neural Network Surrogate Model for the Screening of Therapeutic Antibody Viscosity. Computational and Structural Biotechnology Journal 2022, 20, 2143–2152. 10.1016/j.csbj.2022.04.035.

(31) Olsen, T. H.; Boyles, F.; Deane, C. M. Observed Antibody Space: A Diverse Database of Cleaned, Annotated, and Translated Unpaired and Paired Antibody Sequences. Protein Sci 2022, 31 (1), 141–146. 10.1002/pro.4205.

(32) Dunbar, J.; Deane, C. M. ANARCI: Antigen Receptor Numbering and Receptor Classification. Bioinformatics 2016, 32 (2), 298–300. 10.1093/bioinformatics/btv552.

(33) Dunbar, J.; Krawczyk, K.; Leem, J.; Marks, C.; Nowak, J.; Regep, C.; Georges, G.; Kelm, S.; Popovic, B.; Deane, C. M. SAbPred: A Structure-Based Antibody Prediction Server. Nucleic Acids Research 2016, 44 (W1), W474–W478. 10.1093/nar/gkw361.

(34) Jorgensen, W. L.; Chandrasekhar, J.; Madura, J. D.; Impey, R. W.; Klein, M. L. Comparison of Simple Potential Functions for Simulating Liquid Water. The Journal of Chemical Physics 1983, 79 (2), 926–935. 10.1063/1.445869.

(35) Humphrey, W.; Dalke, A.; Schulten, K. VMD: Visual Molecular Dynamics. J Mol Graph 1996, 14 (1), 33–38, 27–28. 10.1016/0263-7855(96)00018-5.

(36) Klauda, J. B.; Venable, R. M.; Freites, J. A.; O’Connor, J. W.; Tobias, D. J.; Mondragon-Ramirez, C.; Vorobyov, I.; MacKerell, A. D.; Pastor, R. W. Update of the CHARMM All-Atom Additive Force Field for Lipids: Validation on Six Lipid Types. J Phys Chem B 2010, 114 (23), 7830–7843. 10.1021/jp101759q.

(37) Huang, J.; Rauscher, S.; Nawrocki, G.; Ran, T.; Feig, M.; de Groot, B. L.; Grubmüller, H.; MacKerell, A. D. CHARMM36m: An Improved Force Field for Folded and Intrinsically Disordered Proteins. Nat Methods 2017, 14 (1), 71–73. 10.1038/nmeth.4067.

(38) Phillips, J. C.; Braun, R.; Wang, W.; Gumbart, J.; Tajkhorshid, E.; Villa, E.; Chipot, C.; Skeel, R. D.; Kalé, L.; Schulten, K. Scalable Molecular Dynamics with NAMD. Journal of Computational Chemistry 2005, 26 (16), 1781–1802. 10.1002/jcc.20289.

(39) Essmann, U.; Perera, L.; Berkowitz, M. L.; Darden, T.; Lee, H.; Pedersen, L. G. A Smooth Particle Mesh Ewald Method. The Journal of Chemical Physics 1995, 103 (19), 8577–8593. 10.1063/1.470117.

(40) Pedregosa, F.; Varoquaux, G.; Gramfort, A.; Michel, V.; Thirion, B.; Grisel, O.; Blondel, M.; Prettenhofer, P.; Weiss, R.; Dubourg, V.; Vanderplas, J.; Passos, A.; Cournapeau, D. Scikit-Learn: Machine Learning in Python. MACHINE LEARNING IN PYTHON.

(41) Chollet F. Keras. 2015. https://keras.io/ -Google Search. https://www.google.com/search?q=41)+Chollet+F.+Keras.+2015.+https%3A%2F%2Fkeras.io%2F&rlz=1C1GCEB_enUS1031US1031&oq=41)%09Chollet+F.+Keras.+2015.+https%3A%2F%2Fkeras.io%2F&gs_lcrp=EgZjaHJvbWUyBggAEEUYOTIKCAEQABiABBiiBNIBBzUxNmowajSoAgCwAgA&sourceid=chrome&ie=UTF-8 (accessed 2024-02-20).

(42) Abadi, M.; Barham, P.; Chen, J.; Chen, Z.; Davis, A.; Dean, J.; Devin, M.; Ghemawat, S.; Irving, G.; Isard, M.; Kudlur, M.; Levenberg, J.; Monga, R.; Moore, S.; Murray, D. G.; Steiner, B.; Tucker, P.; Vasudevan, V.; Warden, P.; Wicke, M.; Yu, Y.; Zheng, X. TensorFlow: A System for Large-Scale Machine Learning. arXiv May 31, 2016. 10.48550/arXiv.1605.08695.

(43) Team, K. Keras documentation: Getting started with KerasTuner. https://keras.io/guides/keras_tuner/getting_started/ (accessed 2024-02-20).

(44) Dudko, L.; Volkov, P.; Vorotnikov, G.; Zaborenko, A. Application of Deep Learning Technique to an Analysis of Hard Scattering Processes at Colliders. arXiv September 14, 2021. 10.48550/arXiv.2109.08520.

(45) Dietterich, T. Overfitting and Undercomputing in Machine Learning. ACM Comput. Surv. 1995, 27 (3), 326–327. 10.1145/212094.212114.

(46) Raschka, S. MLxtend: Providing Machine Learning and Data Science Utilities and Extensions to Python’s Scientific Computing Stack. Journal of Open Source Software 2018, 3 (24), 638. 10.21105/joss.00638.

(47) Chothia, C.; Lesk, A. M. The Relation between the Divergence of Sequence and Structure in Proteins. EMBO J 1986, 5 (4), 823–826. 10.1002/j.1460-2075.1986.tb04288.x.

(48) Antibody numbering schemes and CDR definitions. https://pipebio.com/blog/antibody-numbering (accessed 2024-02-20).

