## Supporting Information for "DeepSP: Deep Learning-Based Spatial Properties to Predict Monoclonal Antibody Stability"

### Supplementary Data

Table S1: Best Hyperparameters for Approach 1 where individual CNN models were trained for each of the 30 properties.

| | CONV1D_1 | | | CONV1D_2 | | | CONV1D_3 | | | | |
|---|---|---|---|---|---|---|---|---|---|---|---|
| Property | **filter_1** | **kernel_1** | **dropout_1** | **filter_2** | **kernel_2** | **dropout_2** | **filter_3** | **kernel_3** | **dropout_3** | **dense_1** | **learning_rate** |
| SAP_pos_CDRH1 | 96 | 5 | 0.3 | 96 | 4 | 0.3 | 20 | 5 | 0.2 | 32 | 0.001 |
| SAP_pos_CDRH2 | 80 | 5 | 0.2 | 48 | 3 | 0.2 | 32 | 5 | 0.2 | 80 | 0.005 |
| SAP_pos_CDRH3 | 112 | 5 | 0.4 | 48 | 5 | 0.3 | 36 | 5 | 0.3 | 112 | 0.001 |
| SAP_pos_CDRL1 | 128 | 5 | 0.3 | 128 | 5 | 0.4 | 16 | 4 | 0.4 | 32 | 0.001 |
| SAP_pos_CDRL2 | 128 | 3 | 0.2 | 80 | 5 | 0.3 | 76 | 5 | 0.2 | 96 | 0.005 |
| SAP_pos_CDRL3 | 64 | 4 | 0.4 | 112 | 4 | 0.4 | 108 | 5 | 0.2 | 64 | 0.001 |
| SAP_pos_CDR | 96 | 3 | 0.4 | 48 | 3 | 0.2 | 56 | 3 | 0.2 | 32 | 0.005 |
| SAP_pos_Hv | 80 | 4 | 0.4 | 64 | 3 | 0.3 | 84 | 5 | 0.2 | 112 | 0.01 |
| SAP_pos_Lv | 96 | 4 | 0.2 | 128 | 4 | 0.4 | 88 | 4 | 0.2 | 48 | 0.005 |
| SAP_pos_Fv | 128 | 3 | 0.2 | 32 | 3 | 0.2 | 60 | 5 | 0.2 | 96 | 0.01 |
| SCM_pos_CDRH1 | 96 | 4 | 0.3 | 48 | 4 | 0.3 | 76 | 4 | 0.3 | 128 | 0.01 |
| SCM_pos_CDRH2 | 96 | 3 | 0.3 | 96 | 4 | 0.2 | 16 | 3 | 0.2 | 112 | 0.001 |
| SCM_pos_CDRH3 | 80 | 5 | 0.3 | 96 | 5 | 0.4 | 112 | 3 | 0.4 | 128 | 0.005 |
| SCM_pos_CDRL1 | 128 | 3 | 0.2 | 48 | 4 | 0.3 | 68 | 3 | 0.4 | 128 | 0.01 |
| SCM_pos_CDRL2 | 96 | 5 | 0.2 | 64 | 4 | 0.2 | 44 | 5 | 0.2 | 112 | 0.001 |
| SCM_pos_CDRL3 | 128 | 5 | 0.2 | 128 | 5 | 0.3 | 20 | 3 | 0.3 | 48 | 0.005 |
| SCM_pos_CDR | 64 | 3 | 0.2 | 96 | 5 | 0.2 | 116 | 5 | 0.2 | 64 | 0.005 |
| SCM_pos_Hv | 96 | 3 | 0.3 | 96 | 3 | 0.2 | 116 | 5 | 0.2 | 32 | 0.005 |
| SCM_pos_Lv | 80 | 3 | 0.2 | 112 | 3 | 0.4 | 76 | 5 | 0.2 | 64 | 0.005 |
| SCM_pos_Fv | 48 | 3 | 0.3 | 128 | 4 | 0.3 | 100 | 5 | 0.2 | 128 | 0.01 |
| SCM_neg_CDRH1 | 48 | 4 | 0.2 | 48 | 3 | 0.2 | 124 | 4 | 0.2 | 128 | 0.001 |
| SCM_neg_CDRH2 | 32 | 3 | 0.4 | 48 | 5 | 0.2 | 124 | 5 | 0.2 | 112 | 0.001 |
| SCM_neg_CDRH3 | 48 | 5 | 0.3 | 80 | 5 | 0.3 | 68 | 5 | 0.2 | 64 | 0.005 |
| SCM_neg_CDRL1 | 80 | 5 | 0.3 | 112 | 4 | 0.3 | 68 | 4 | 0.2 | 32 | 0.005 |
| SCM_neg_CDRL2 | 112 | 4 | 0.3 | 16 | 4 | 0.2 | 80 | 5 | 0.2 | 80 | 0.005 |
| SCM_neg_CDRL3 | 112 | 5 | 0.2 | 80 | 4 | 0.2 | 80 | 4 | 0.3 | 48 | 0.005 |
| SCM_neg_CDR | 96 | 4 | 0.2 | 96 | 5 | 0.3 | 96 | 3 | 0.2 | 96 | 0.01 |
| SCM_neg_Hv | 96 | 3 | 0.4 | 128 | 5 | 0.3 | 20 | 5 | 0.2 | 64 | 0.01 |
| SCM_neg_Lv | 48 | 4 | 0.3 | 112 | 5 | 0.3 | 40 | 4 | 0.2 | 96 | 0.005 |
| SCM_neg_Fv | 128 | 5 | 0.2 | 16 | 3 | 0.2 | 48 | 5 | 0.2 | 128 | 0.01 |

Table S2: Model performance metrics for Approach 1 where individual CNN models were trained for each of the 30 properties.

| Property | Mean_score | Baseline_MAE | Val_loss | MAE | R |
|---|---|---|---|---|---|
| **SAP_pos_CDRH1** | 2.63 | 1.42 | 0.68 | **0.70** | **0.82** |
| **SAP_pos_CDRH2** | 2.74 | 2.31 | 0.58 | **0.56** | **0.97** |
| **SAP_pos_CDRH3** | 14.90 | 7.05 | 2.39 | **2.41** | **0.93** |
| **SAP_pos_CDRL1** | 3.17 | 2.12 | 0.74 | **0.75** | **0.91** |
| **SAP_pos_CDRL2** | 2.39 | 1.50 | 0.44 | **0.47** | **0.94** |
| **SAP_pos_CDRL3** | 5.51 | 2.59 | 1.42 | **1.43** | **0.77** |
| **SAP_pos_CDR** | 31.34 | 8.37 | 3.72 | **3.70** | **0.87** |
| **SAP_pos_Hv** | 58.04 | 8.47 | 4.15 | **4.09** | **0.84** |
| **SAP_pos_Lv** | 42.14 | 7.14 | 3.26 | **3.36** | **0.80** |
| **SAP_pos_Fv** | 100.18 | 11.55 | 6.05 | **6.28** | **0.73** |
| **SCM_pos_CDRH1** | 47.45 | 25.95 | 12.26 | **12.20** | **0.86** |
| **SCM_pos_CDRH2** | 29.60 | 23.88 | 7.83 | **8.14** | **0.90** |
| **SCM_pos_CDRH3** | 76.95 | 50.61 | 23.24 | **24.18** | **0.86** |
| **SCM_pos_CDRL1** | 68.01 | 34.16 | 13.52 | **13.24** | **0.91** |
| **SCM_pos_CDRL2** | 63.97 | 29.37 | 10.30 | **10.45** | **0.92** |
| **SCM_pos_CDRL3** | 47.23 | 30.09 | 14.91 | **14.88** | **0.82** |
| **SCM_pos_CDR** | 333.21 | 111.87 | 48.29 | **49.17** | **0.89** |
| **SCM_pos_Hv** | 1178.35 | 166.39 | 81.25 | **81.47** | **0.86** |
| **SCM_pos_Lv** | 1045.27 | 129.72 | 61.60 | **60.87** | **0.88** |
| **SCM_pos_Fv** | 2223.61 | 221.47 | 108.18 | **105.63** | **0.87** |
| **SCM_neg_CDRH1** | 44.01 | 28.19 | 12.10 | **12.46** | **0.91** |
| **SCM_neg_CDRH2** | 37.17 | 28.83 | 9.01 | **9.05** | **0.94** |
| **SCM_neg_CDRH3** | 116.38 | 67.77 | 31.15 | **30.79** | **0.88** |
| **SCM_neg_CDRL1** | 67.22 | 33.99 | 13.59 | **13.58** | **0.94** |
| **SCM_neg_CDRL2** | 27.60 | 20.41 | 6.82 | **6.89** | **0.93** |
| **SCM_neg_CDRL3** | 86.19 | 37.57 | 20.84 | **20.55** | **0.81** |
| **SCM_neg_CDR** | 378.56 | 134.63 | 59.26 | **57.92** | **0.91** |
| **SCM_neg_Hv** | 498.95 | 147.78 | 70.07 | **68.60** | **0.89** |
| **SCM_neg_Lv** | 443.74 | 114.18 | 46.46 | **46.72** | **0.92** |
| **SCM_neg_Fv** | 942.69 | 206.09 | 91.98 | **90.05** | **0.90** |

Table S3: Best Hyperparameters for Approach 2, where 3 models were trained for each property to predict in 10 domains of antibodies. [10 domains – CDRH1, CDRH2, CDRH3, CDRL1, CDRL2, CDRL3, CDR, Hv, Lv, Fv]

| property | filter_1 | kernel_1 | dropout_1 | filter_2 | kernel_2 | dropout_2 | filter_3 | kernel_3 | dense_1 | Dense_2 | learning_rate | Output Layer |
|---|---|---|---|---|---|---|---|---|---|---|---|---|
| SAP_pos | 128 | 5 | 0.3 | 96 | 4 | - | 32 | 5 | 112 | 48 | 0.0001 | 10 |
| SCM_pos | 128 | 4 | 0.4 | 112 | 4 | 0.4 | 144 | 5 | 128 | - | 0.005 | 10 |
| SCM_neg | 128 | 5 | 0.1 | 112 | 4 | - | 64 | 4 | 128 | - | 0.0001 | 10 |

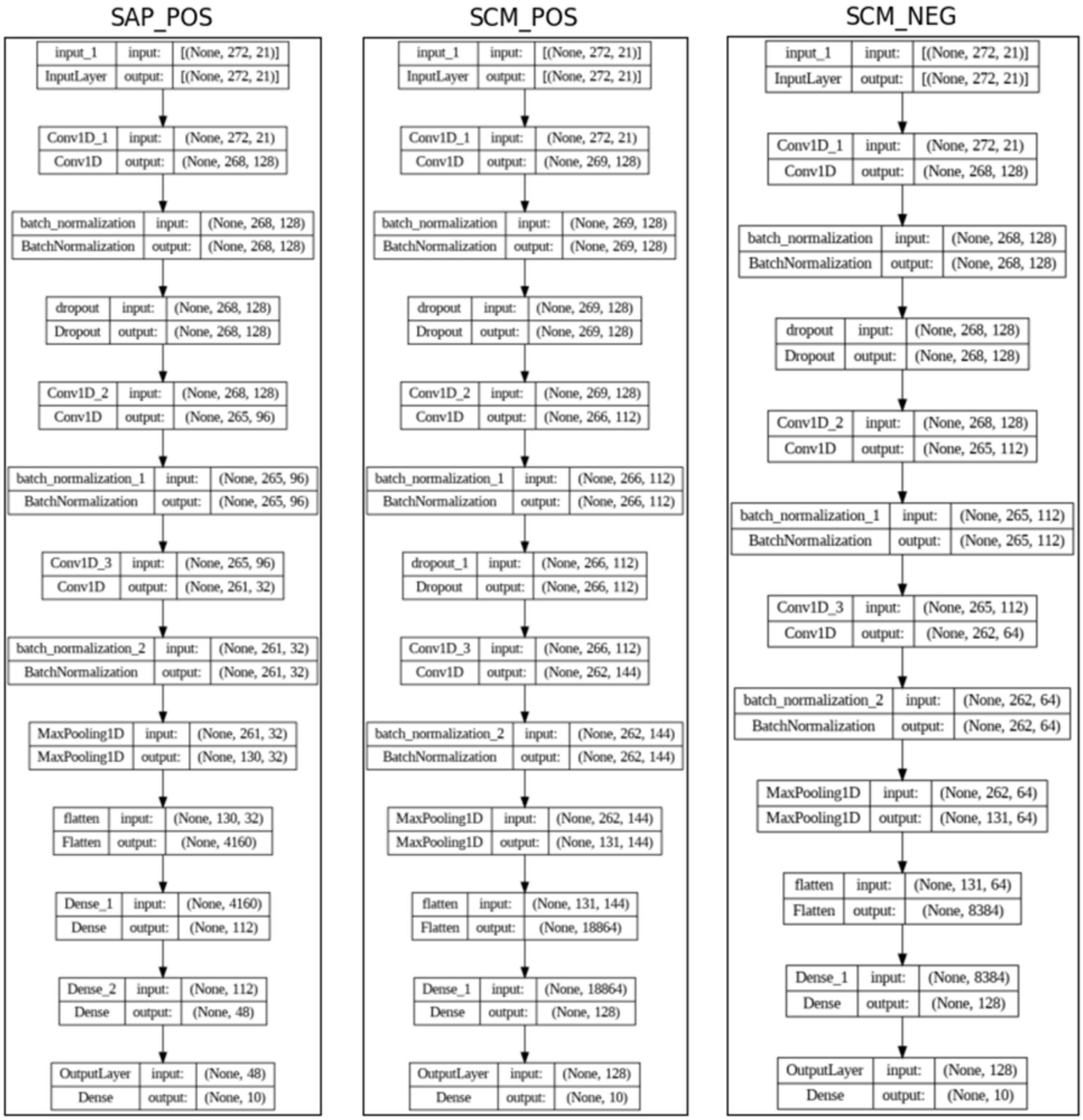

Table S4: Model performance metrics for Approach 2 where 3 models were trained for each property to predict in 10 domains of antibodies. [10 domains – CDRH1, CDRH2, CDRH3, CDRL1, CDRL2, CDRL3, CDR, Hv, Lv, Fv]

| Property | Mean_score | Baseline_MAE | Val_loss | MAE | R |
|---|---|---|---|---|---|
| SAP_pos_CDRH1 | 2.63971335 | 1.378985788 | 2.35556 | 0.7541 | 0.80408 |
| SAP_pos_CDRH2 | 2.81707453 | 2.40223994 | 2.35556 | 0.616 | 0.96342 |
| SAP_pos_CDRH3 | 14.5431266 | 6.826758637 | 2.35556 | 2.3927 | 0.92934 |
| SAP_pos_CDRL1 | 3.13093911 | 2.07825485 | 2.35556 | 0.7945 | 0.89581 |
| SAP_pos_CDRL2 | 2.42763906 | 1.538941623 | 2.35556 | 0.6592 | 0.91611 |
| SAP_pos_CDRL3 | 5.54790209 | 2.663020266 | 2.35556 | 1.4355 | 0.76548 |
| SAP_pos_CDR | 31.1075857 | 8.225197621 | 2.35556 | 3.7066 | 0.86873 |
| SAP_pos_Hv | 57.7602197 | 8.206724212 | 2.35556 | 4.1181 | 0.83011 |
| SAP_pos_Lv | 42.2474328 | 7.294081579 | 2.35556 | 3.4564 | 0.79301 |
| SAP_pos_Fv | 100.00781 | 11.28239503 | 2.35556 | 6.2045 | 0.73277 |
| SCM_pos_CDRH1 | 46.2467075 | 26.11127996 | 38.6219 | 13.714 | 0.8403 |
| SCM_pos_CDRH2 | 29.2702336 | 23.72153664 | 38.6219 | 9.4191 | 0.89753 |
| SCM_pos_CDRH3 | 76.1268933 | 50.81273015 | 38.6219 | 25.105 | 0.85688 |
| SCM_pos_CDRL1 | 67.8014089 | 34.68469895 | 38.6219 | 14.138 | 0.89631 |
| SCM_pos_CDRL2 | 62.9204742 | 28.90342605 | 38.6219 | 13.477 | 0.91035 |
| SCM_pos_CDRL3 | 45.7080582 | 28.82579346 | 38.6219 | 16.003 | 0.79936 |
| SCM_pos_CDR | 328.077582 | 111.6501535 | 38.6219 | 48.695 | 0.88946 |
| SCM_pos_Hv | 1171.43712 | 163.1671595 | 38.6219 | 84.456 | 0.85475 |
| SCM_pos_Lv | 1043.1453 | 128.1051035 | 38.6219 | 58.952 | 0.88026 |
| SCM_pos_Fv | 2214.582 | 219.9718012 | 38.6219 | 103.05 | 0.8766 |
| SCM_neg_CDRH1 | 46.7682409 | 29.80162768 | 36.2933 | 12.735 | 0.90447 |
| SCM_neg_CDRH2 | 38.55516 | 30.07687559 | 36.2933 | 9.8978 | 0.93604 |
| SCM_neg_CDRH3 | 118.35354 | 69.85537506 | 36.2933 | 32.798 | 0.86935 |
| SCM_neg_CDRL1 | 68.165604 | 35.78393596 | 36.2933 | 13.877 | 0.93705 |
| SCM_neg_CDRL2 | 27.7998108 | 20.36697021 | 36.2933 | 7.4208 | 0.92127 |
| SCM_neg_CDRL3 | 86.8210701 | 37.03727339 | 36.2933 | 20.972 | 0.80537 |
| SCM_neg_CDR | 386.457136 | 140.2793733 | 36.2933 | 57.297 | 0.90987 |
| SCM_neg_Hv | 510.159462 | 150.8213874 | 36.2933 | 70.471 | 0.889 |
| SCM_neg_Lv | 445.946364 | 117.0513646 | 36.2933 | 46.274 | 0.92257 |
| SCM_neg_Fv | 956.103832 | 210.1025702 | 36.2933 | 90.054 | 0.90454 |

Table S5: Summary of Spatial Properties Analysis in Different Antibody Regions Obtained from MD Simulations.

| Property | Minimum | Maximum | Q1 | Q2 | Q3 | NumBelow LowerWhisker | NumAbove UpperWhisker |
|---|---|---|---|---|---|---|---|
| SAPposCDRH1 | 0.006 | 25.166 | 1.428 | 2.06 | 2.938 | 0 | 2052 |
| SAPposCDRH2 | 0 | 28.11 | 0.276 | 1.852 | 3.5915 | 0 | 1369 |
| SAPposCDRH3 | 0 | 66.986 | 8.2125 | 13.39 | 20.0055 | 0 | 358 |
| SAPposCDRL1 | 0 | 24.472 | 1.222 | 2.446 | 4.412 | 0 | 700 |
| SAPposCDRL2 | 0 | 20.334 | 0.996 | 1.678 | 3.2255 | 0 | 855 |
| SAPposCDRL3 | 0.04 | 33.608 | 3.118 | 4.758 | 7.204 | 0 | 675 |
| SAPposCDR | 3.39 | 100.542 | 23.68 | 30.142 | 37.648 | 0 | 306 |
| SAPposHv | 25.526 | 130.676 | 50.536 | 57.018 | 64.4835 | 9 | 330 |
| SAPposLv | 21.558 | 104.372 | 35.34 | 40.675 | 47.292 | 0 | 453 |
| SAPpos_Fv | 58.3 | 218.262 | 89.944 | 98.528 | 108.335 | 4 | 555 |
| SCMposCDRH1 | 0.018 | 255.824 | 22.806 | 40.631 | 65.1895 | 0 | 483 |
| SCMposCDRH2 | 0 | 325.324 | 7.574 | 17.24 | 44.191 | 0 | 800 |
| SCMposCDRH3 | 0.368 | 702.722 | 28.204 | 55.857 | 104.0775 | 0 | 974 |
| SCMposCDRL1 | 1.156 | 474.062 | 35.893 | 57.193 | 91.615 | 0 | 532 |
| SCMposCDRL2 | 0.696 | 295.694 | 34.543 | 60.297 | 84.531 | 0 | 389 |
| SCMposCDRL3 | 0.284 | 433.476 | 19.152 | 33.474 | 62.225 | 0 | 1070 |
| SCMposCDR | 16.784 | 1402.992 | 229.494 | 310.355 | 413.477 | 0 | 397 |
| SCMposHv | 467.284 | 2231.214 | 1032.19 | 1168.81 | 1311.391 | 40 | 198 |
| SCMposLv | 371.906 | 1946.634 | 939.496 | 1041.44 | 1146.045 | 103 | 295 |
| SCMposFv | 1230.422 | 3700.66 | 2030.94 | 2210.71 | 2401.32 | 56 | 182 |
| SCMnegCDRH1 | 0.404 | 341.812 | 17.809 | 32.257 | 56.605 | 0 | 1244 |
| SCMnegCDRH2 | 0 | 364.096 | 9.3375 | 23.697 | 53.8145 | 0 | 798 |
| SCMnegCDRH3 | 0.068 | 753.508 | 50.3025 | 96.187 | 160.683 | 0 | 598 |
| SCMnegCDRL1 | 1.322 | 443.018 | 36.7605 | 52.723 | 79.6135 | 0 | 1334 |
| SCMnegCDRL2 | 0.058 | 222.816 | 8.6745 | 17.765 | 35.14 | 0 | 1770 |
| SCMnegCDRL3 | 0.898 | 404.236 | 51.526 | 78.574 | 111.6285 | 0 | 577 |
| SCMnegCDR | 41.576 | 1567.17 | 251.69 | 348.292 | 472.94 | 0 | 517 |
| SCMnegHv | 91.81 | 2056.878 | 361.19 | 467.681 | 601.9535 | 0 | 522 |
| SCMnegLv | 118.286 | 1414.12 | 339.52 | 410.924 | 506.265 | 0 | 1001 |
| SCMnegFv | 275.898 | 2587.318 | 753.26 | 906.613 | 1092.288 | 0 | 466 |

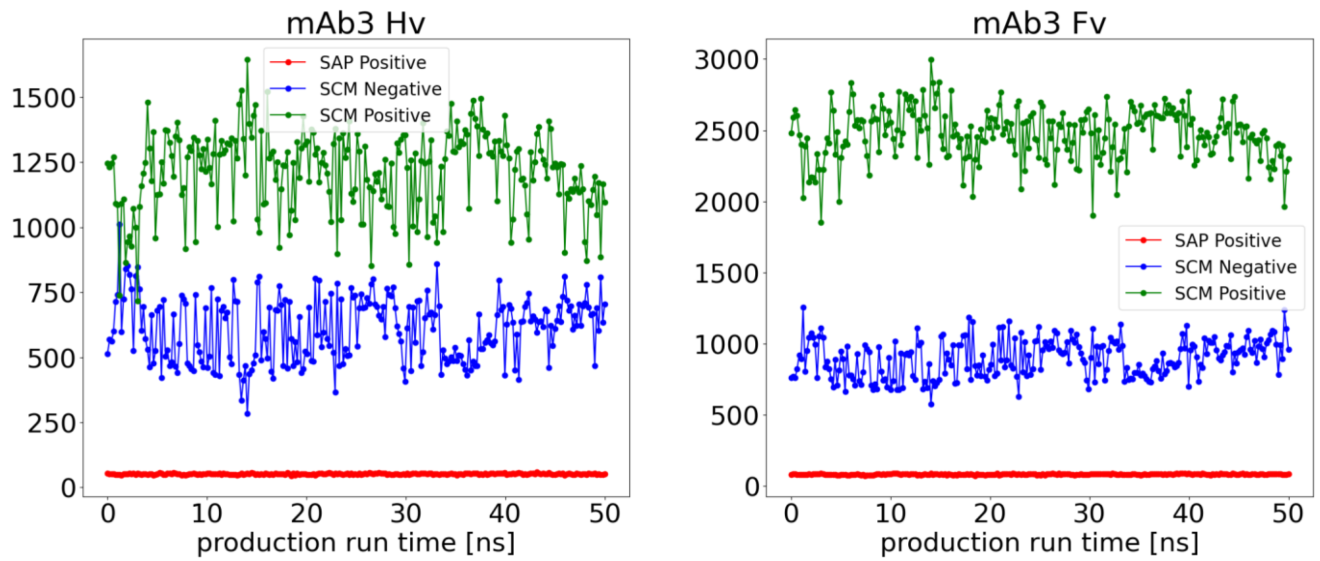

Figure S1: Variations in SAP positive, SCM negative and SCM positive scores in the variable region of the heavy chain (Hv) and the entire variable region (Fv) of an antibody.

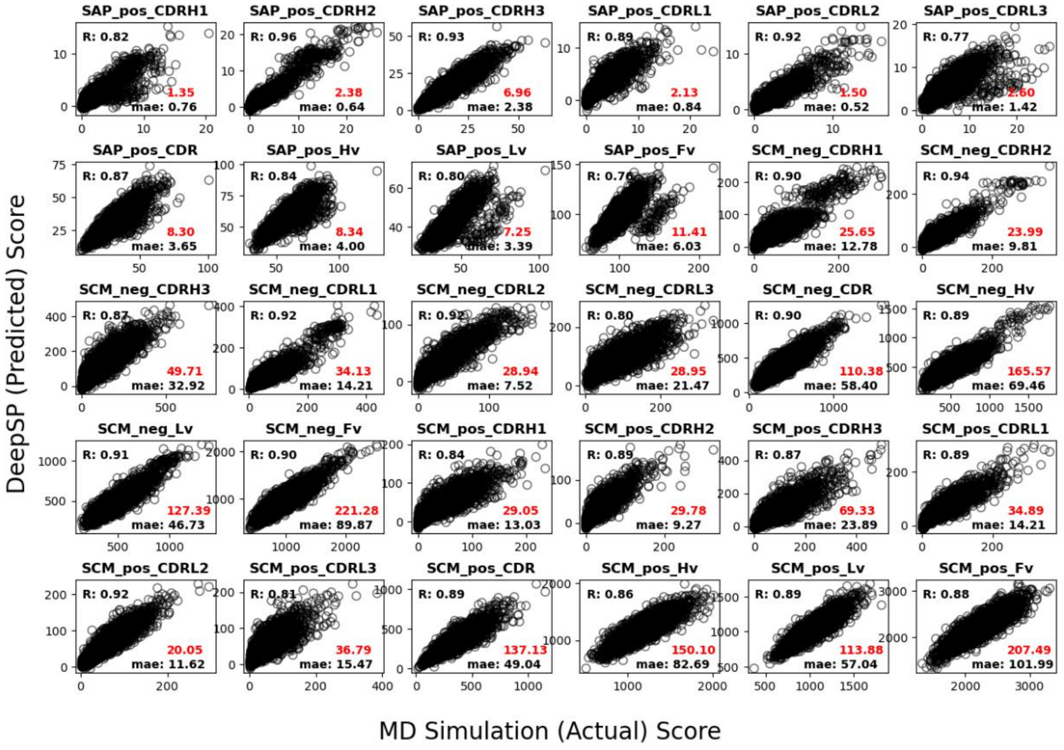

Figure S2: Scatter plot illustrating the correlation between the predicted and actual score, and MAE of all 30 spatial properties. (baseline MAE is shown in red).

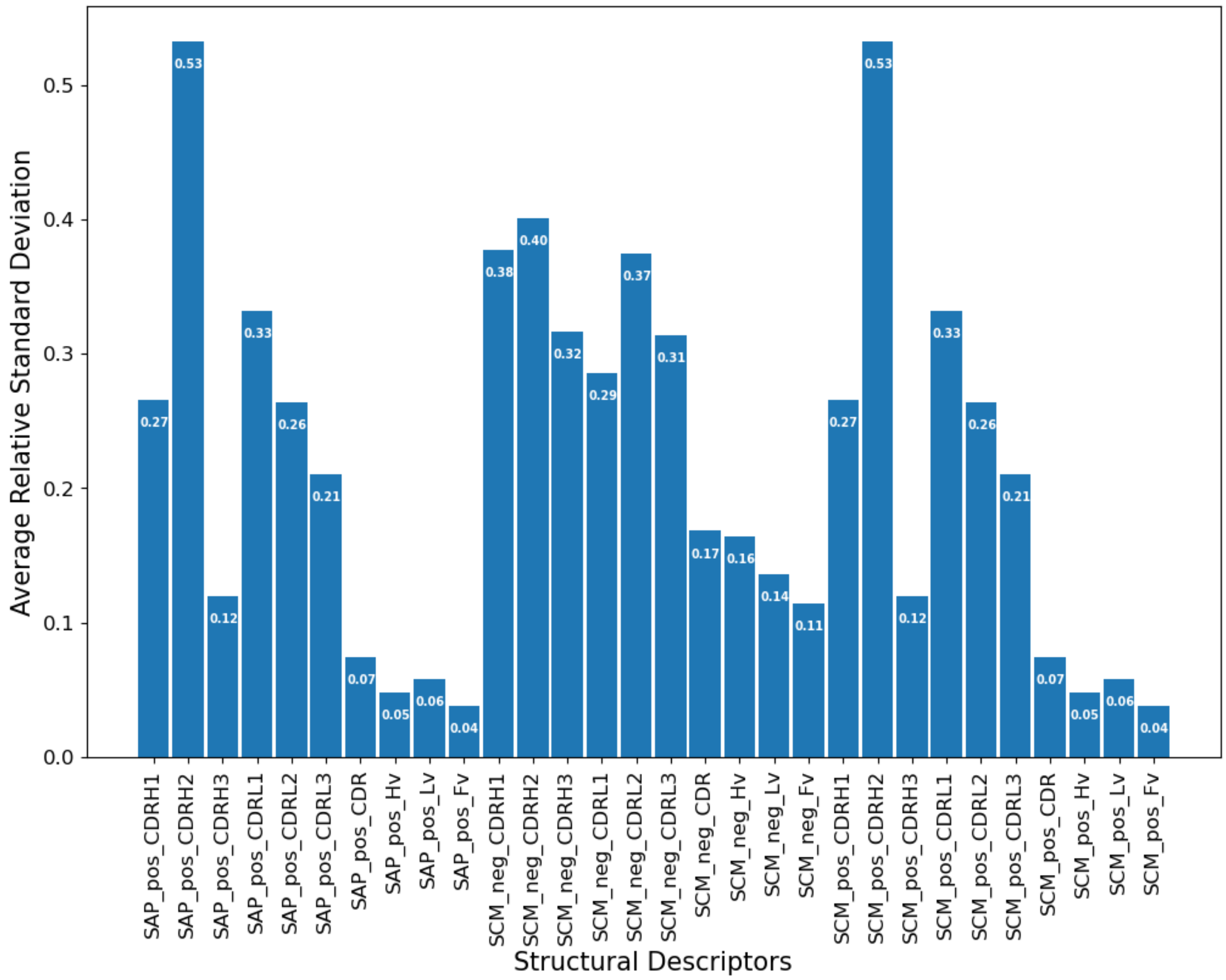

Figure S3. Average Relative Standard deviation of all 30 spatial properties. The relative standard deviation was obtained by dividing the standard deviation by the actual average value.

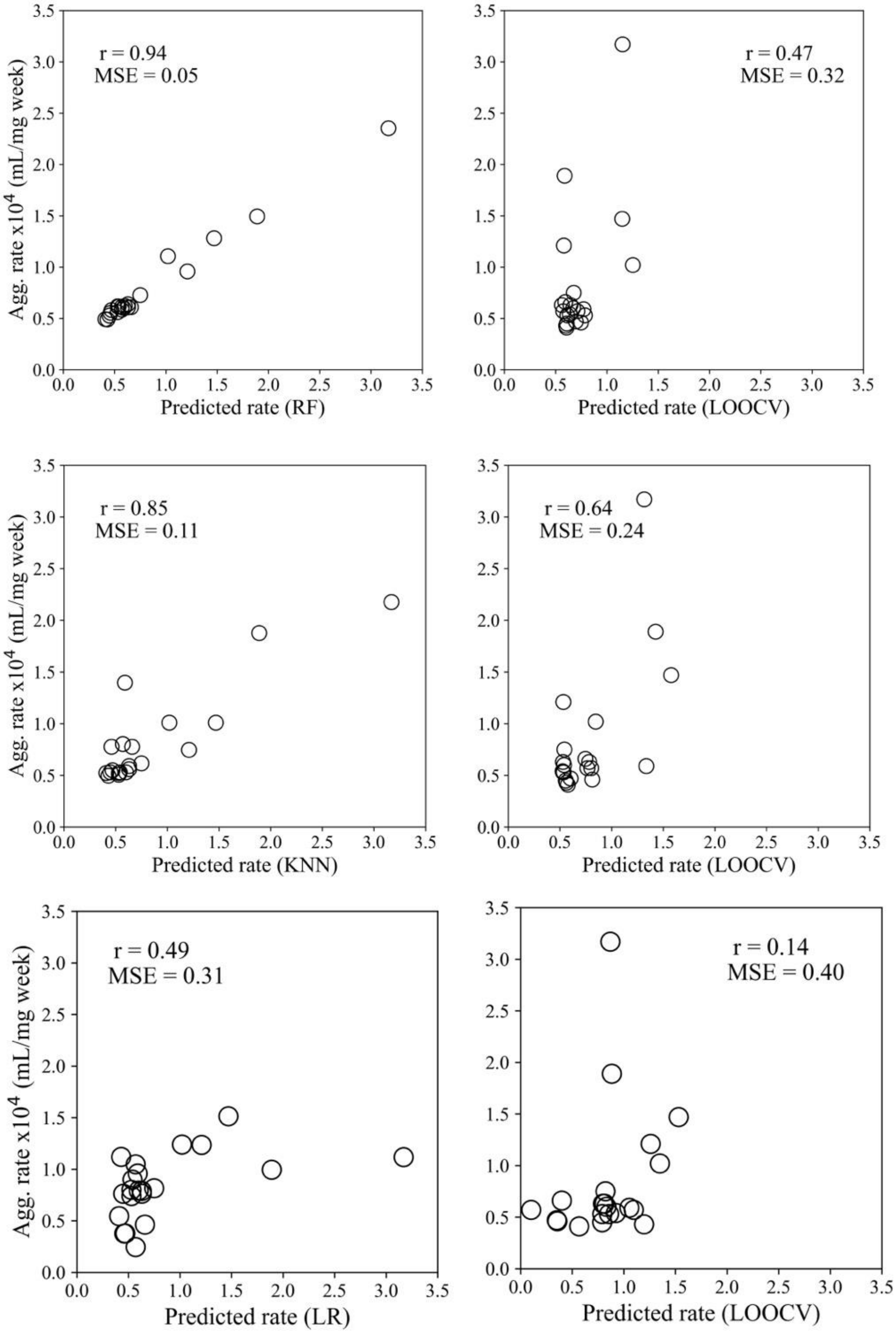

Figure S4. Correlation coefficients for the 3-feature or 4-feature random forest regression, nearest neighbors' regression, and linear regression (LR) models, trained using the entire dataset of 21 samples and LOOCV.
